# The *Neosartorya (Aspergillus) fischeri* antifungal protein NFAP2 has low potential to trigger resistance development in *Candida albicans in vitro*

**DOI:** 10.1101/2024.05.14.594093

**Authors:** Gábor Bende, Nóra Zsindely, Krisztián Laczi, Zsolt Kristóffy, Csaba Papp, Attila Farkas, Liliána Tóth, Szabolcs Sáringer, László Bodai, Gábor Rákhely, Florentine Marx, László Galgóczy

## Abstract

Due to the increase in the number of drug-resistant *Candida albicans* strains, new antifungal compounds with limited potential for development of resistance are urgently needed. NFAP2, an antifungal protein (AFP) secreted by *Neosartorya* (*Aspergillus*) *fischeri*, is a promising candidate. We investigated the ability of *C. albicans* to develop resistance to NFAP2 in a microevolution experiment compared with generic fluconazole (FLC). *C. albicans* adapted to only 1 × minimum inhibitory concentration (MIC) of NFAP2 compared with 32 × MIC of FLC. Genome analysis revealed non-silent mutations in only two genes in NFAP2-resistant strains and in several genes in FLC-resistant strains. Resistance development to NFAP2 did not influence cell morphology. The susceptibility of NFAP2-resistant strains did not change to FLC, amphotericin B, micafungin, terbinafine. These strains did not show altered susceptibility to AFPs from *Penicillium chrysogenum*, except one which had less susceptibility to *P. chrysogenum* antifungal protein B. FLC-resistant strains had decreased susceptibility to terbinafine and NFAP2, but not to other drugs and AFPs from *P. chrysogenum*. NFAP2- and FLC-resistant strains showed decreased and increased NFAP2 binding and uptake, respectively. The development of resistance to NFAP2 decreased tolerance to cell wall, heat, and UV stresses. The development of FLC resistance increased tolerance to cell wall stress and decreased tolerance to heat and UV stresses. Resistance to NFAP2 did not have significant metabolic fitness cost and could not increase virulence, compared with resistance to FLC.

**Importance:** Due to the increasing number of (multi)drug-resistant strains, only a few effective antifungal drugs are available to treat infections caused by opportunistic *Candida* species. Therefore, the incidence of hard-to-treat candidiasis has increased dramatically in the past decade, and the demand to identify antifungal compounds with minimal potential to trigger resistance is substantial. The features of NFAP2 make it a promising candidate for the topical treatment of *Candida* infection. Data on the development of resistance to AFPs in *C. albicans* are lacking. In this study, we provide evidence that NFAP2 has low potential to trigger resistance in *C. albicans in vitro* and the developed resistance mechanisms to NFAP2 are not associated with severe phenotypic changes compared with development of resistance to generic FLC. These results suggest the slow emergence of NFAP2-resistant *Candida* strains and that NFAP2 can reliably be used long-term in the clinic.

## Introduction

In the last two decades, the occurrence of opportunistic fungal infections (FIs) has increased due the the rise in the number of patients with immunosuppression and resistance to antifungal drugs. This increase poses a serious threat to public health globally (Poissy et al.,2022). In addition to causing nonlife-threatening cutaneous mycoses in immunocompetent individuals, fungi cause subcutaneous or systemic and disseminated fatal infections in patients with a weak and impaired immune system due to HIV infection, hematologic malignancies, diabetes mellitus, immunosuppressive therapy, or intensive care of organ transplantation (Friedman et al., 2019; Hossain et al., 2022). FIs have generated little public attention compared with tuberculosis and malaria, although the number of annual deaths (∼1.7 million) due to FIs is close to the number of deaths due to tuberculosis and malaria together (Kainz et al., 2020). Therefore, the World Health Organization released the first fungal priority pathogens list in October 2022, which highlights 19 fungal pathogens that pose the most serious threat to human health. Five of the listed fungi belong to the genus *Candida* (World Health Organization, 2022).

Candidiasis, caused by *Candida* species, is the most common FI (Bongomin et al., 2017) and represents one of the most prevalent hospital-acquired infections (Rautemaa-Richardson et al., 2017). *Candida albicans* is responsible for most cases; however, an epidemiological shift to non-*albicans Candida* (NAC) species, represented by *Nakaseomyces glabrata* (*Candida glabrata*), *Candida tropicalis, Candida parapsilosis*, and *Pichia kudriavzevii* (*Candida krusei*), has been observed recently. Depending on the age group and geographical location of the patient, the NAC species can exceed *C. albicans* (Pappas et al., 2016). The emergence of multidrug-resistant nosocomial *Candida auris* is a cause of alarm in epidemiology (Friedman et al., 2019). Treatment of *Candida* infection is based on a relatively shallow pool of four antifungal drug classes: azoles, echinocandins, polyenes, and fluorinated pyrimidines (Ksiezopolska et al., 2018). Fluconazole (FLC) is the most commonly used fungistatic agent for treating candidiasis; therefore, it is not surprising that FLC resistance is common among human pathogenic *Candida* species (Perlin et al., 2017). Resistance against a certain class of antifungal drugs severely hampers good therapeutic outcome and requires the administration of other classes of antifungals (Poissy et al., 2022). The increased use of another class of antifungals in combination with a less effective drug increases the risk of multidrug resistance (Ksiezopolska et al., 2018). For example, the introduction of echinocandins to treat *N. glabrata* infections clinically led to the development of multidrug resistance to azoles–echinocandins (Alexander et al., 2013). Emerging (multi)drug-resistance and lack of a new class of antifungal agents with novel mechanisms of action have led to an urgent demand for new antifungal strategies to treat FIs caused not only by *Candida* species, but also other human fungal pathogens.

Several new antifungal drugs are in various stages of clinical trials. They offer new avenues for the treatment of FIs due to their novel modes of action, with reduced toxicity, less adverse effects, and minimal resistance development potential compared with conventional drugs. Among others, rezafungin (CD101), ibrexafungerp (SCY-078), and VT-1161 are the most promising candidates, which are highly active against several *Candida* species, including conventional drug-resistant *C. albicans* and *C. auris* isolates, and have passed some phases of clinical trial (Rauseo et al., 2020). Although several new antifungal compounds are in the early stages of clinical trial, very few have reached late stages, which does not guarantee marketed treatment (Van Daele et al., 2019). Therefore, the search for new antifungal compounds is essential. Biomolecules with antifungal effects are promising alternatives (Augostine et al., 2022).

Antifungal proteins (AFPs) are promising drug candidates for treating mycoses, because their mode of action differs from conventional antifungal medicines. AFPs can provide less toxic treatment options compared with licensed antifungals, which cause serious adverse effects after prolonged therapy (Fernández de Ullivarri et al., 2020). AFPs of filamentous fungal origin effectively inhibit the growth of different *Candida* species *in vitro* (Galgóczy et al., 2019). NFAP2, an AFP derived from the ascomycete *Neosartorya* (*Aspergillus*) *fischeri* NRRL 181, has high therapeutic potential in the treatment of superficial *Candida* infections (Holzknecht et al., 2022; Kovács et al., 2019;). This small molecular weight (5.6 kDa), cationic, and highly stable extracellular protein is remarkably active against not only planktonic cells of different *Candida* spp. (Tóth et al., 2018), but also sessile biofilm cells of fluconazole-resistant *C. albicans* and multidrug-resistant *C. auris* isolates (Kovács et al., 2019, 2021). NFAP2 can interact synergistically with azoles, echinocandi/ns (first-line antifungal drugs for superficial and/or invasive *Candida* infection), and amphotericin B, significantly decreasing their minimum inhibitory concentrations (MIC) *in vitro* (Kovács et al., 2019, 2021; Tóth et al., 2018). In a murine vulvovaginal candidiasis model, NFAP2 demonstrated its monotherapeutic potential against an FLC-resistant strain of *C. albicans.* Coadministration of NFAP2 and FLC during therapy resulted in synergy *in vivo* and reversed resistance; moreover, none of the treatments caused morphological alterations and serious pathological reactions in vaginal and vulvar tissues (Kovács et al., 2019). Consistently, in a three-dimensional human cell-based skin infection model, NFAP2 was a safe and effective topical agent. NFAP2 significantly decreased the fungal burden of *C. albicans* in the stratum corneum and was well-tolerated by the model, because alterations to an intact outside-in permeability barrier were not observed after NFAP2 treatment (Holzknecht et al., 2022).

Therefore, NFAP2 is a promising therapeutic agent for treating superficial drug-resistant *Candida* infections. One of the requirements for the reliable therapeutic application of NFAP2 involves understanding the potential for development of NFAP2 resistance in fungi.

In this study, our objective was to investigate the potential of NFAP2 to trigger resistance development in *C. albicans* compared with FLC, a widely used anti-*Candida* drug, which is known to commonly induce resistance in *Candida* (Perlin et al., 2017). The results of the microevolution experiments showed that *C. albicans* has limited ability to develop strong resistance to NFAP2, compared with FLC. The development of resistance to NFAP2 does not have a fitness cost and does not significantly alter cell morphology and susceptibility to conventional antifungal drugs, except for FLC. However, it affects uptake of NFAP2, tolerance to abiotic stress, and virulence of *C. albicans*.

## Results

### Generation of NFAP2- and FLC-resistant *C. albicans* strains

Microevolution experiments including six-six independent parallels were performed to investigate the adaptation potential of *C. albicans* to increasing concentrations of NFAP2, compared with FLC, a widely used anti-*Candida* drug (Govindarajan et al., 2023). Other microevolution studies have shown that *C. albicans* can adapt to increasing FLC concentration and evolve strong resistance (Morschhäuser, 2016). Therefore, FLC served as a control for the development of adaptation and resistance mechanisms in our study. First, the susceptibilities of *C. albicans* CBS 5982 to NFAP2 and FLC were determined in low-cation medium (LCM) for the microevolution experiments. The MICs of NFAP2 and FLC were 3.125 and 128 µg mL^−1^, respectively.

Adaptation analysis in the microevolution experiment showed that *C. albicans* CBS 5982 could adapt only to 1 × MIC of NFAP2 (3.125 µg mL^−1^) and did not survive serial passage in the presence of 2 × MIC of NFAP2 (Figure 1A). By contrast, *C. albicans* CBS 5982 could grow even at 32 × MIC of FLC (8192 µg mL^−1^) (Figure 1B). These observations indicated that *C. albicans* has lower potential to adapt to NFAP2 than FLC. The cultures from the adaptation experiment underwent serial passages in antifungal-free medium and cultured on selective LCM agar plates containing 1 × MIC of NFAP2 or 32 × MIC of FLC to remove strains that do not have acquired resistance. This last step of the microevolution experiment selected for stably resistant strains of *C. albicans* that evolved resistance to 1 × MIC of NFAP2 or 32 × MIC of FLC. Three colonies each of independently evolved *C. albicans* cultures resistant to NFAP2 and FLC were randomly selected. These are referred to as NFAP2- and FLC-resistant strains in this study, and they were named NFAP2/4, NFAP2/5, and NFAP2/6 (NFAP2-resistant), as well as FLC/3, FLC/5, FLC/6 (FLC-resistant) strains, respectively. Results of the microevolution experiment showed that *C. albicans* has limited ability to develop pronounced resistance to NFAP2 compared with the conventional triazole antifungal FLC.

**Figure 1.**
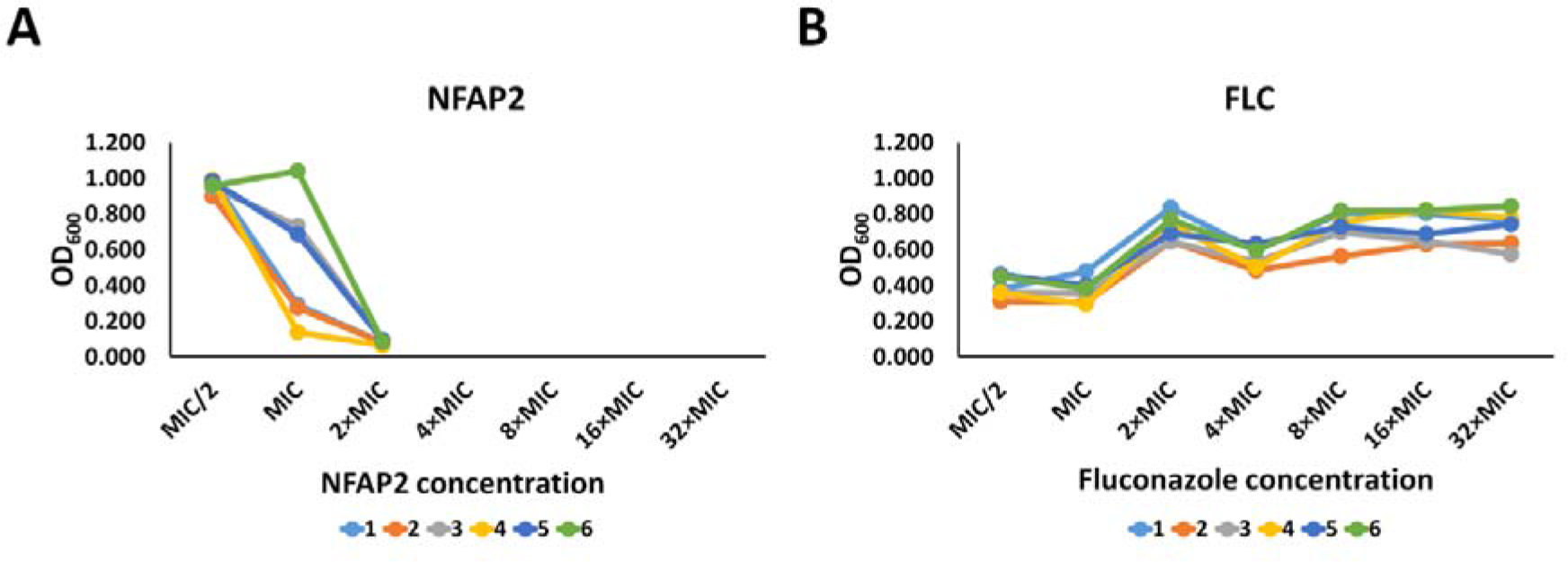
Adaptation of *Candida albicans* CBS 5982 to increasing concentrations of *Neosartorya* (*Aspergillus*) *fischeri* antifungal protein 2 (NFAP2) (A) and fluconazole (FLC) (B) in a microevolution experiment performed in a low-cation medium. The optical densities of cultures of six independent lineages (1–6) were measured at the final passage of each concentration. The minimum inhibitory concentrations (MICs) of NFAP2 and FLC were 3.125 and 128 µg mL^−1^, respectively.

Whole genome sequencing of NFAP2- and FLC-resistant *C. albicans* strains.

The whole genomes of wild-type *C. albicans* CBS 5982 (WT), three independently evolved NFAP2- and FLC-resistant strains were sequenced to reveal genomic changes under NFAP2 or FLC pressure and to identify mutations in resistant lines. Three strains of non-selected control lines that had not been subjected to drug treatment were also sequenced and compared to exclude the influence of the experimental condition on the genome mutation. Considering that *C. albicans* is a diploid yeast, sequences of mutation-affected regions of these genes were checked for heterozygosity and compared with WT sequences, to ensure that these are not artifacts of the genome analysis software. Supplementary Figure 1 shows the sequenograms of the verified mutant gene regions compared with the WT. Whole genome sequencing (WGS) revealed a deletion in a biofilm-induced uncharacterized protein (UBIP; NCBI accession number: XM_709449.2, GenBank: KHC66060.1) in NFAP2/4. NFAP2/5 and NFAP2/6 carried the same multiple nucleotide variation (MNV) at the same position in a putative glycosylphosphatidylinositol-anchored protein PGA58 (NCBI accession number: XM_715546.1). For PGA58, the presence of the heterozygous mutation could not be verified because we obtained noisy and incomplete spectra even after several repetitions of the PCR with different primer pairs (data not shown). FLC/3 contained two single SNVs and one nucleotide insertion in clathrin heavy chain 1 (CHC1; National Center for Biotechnology Information [NCBI] accession number: XM_705744.2,) and two nucleotide deletions in serine/threonine-specific protein phosphatase of type 2C-related family (PTC2; NCBI accession number: XM_713113.2,). FLC/5 had one SNV in phosphoinositide 5-phosphatase (INP51; NCBI accession number: XM_709454.2). FLC/6 contained one SNV in an alkaline-responsive transcriptional regulator RIM101 (NCBI accession number: XM_709954.1). These results are summarized in Table 1. The locations of the mutations in the translated protein sequences are presented in SUPPLEMENTARY TABLE 1 and Supplementary Figure 2.

**Table 1.**
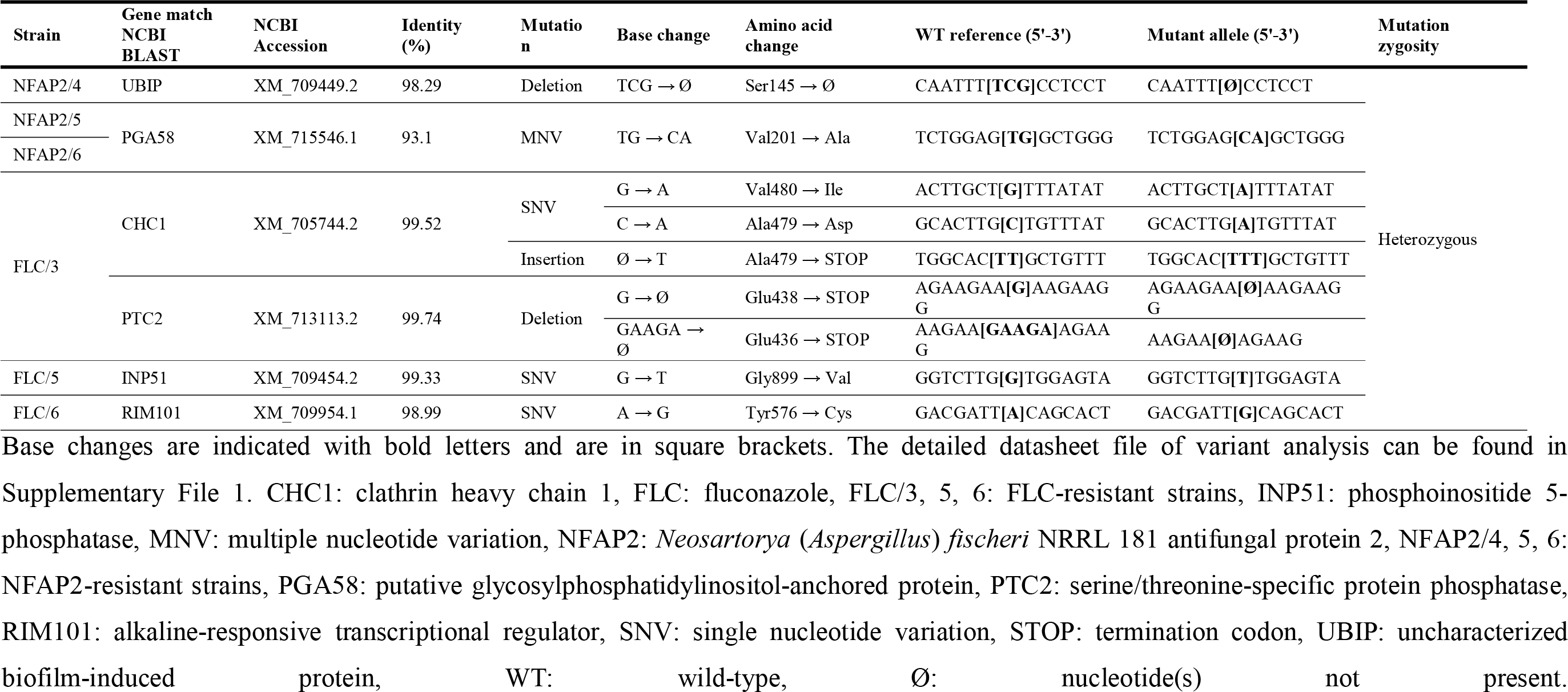
National Center for Biotechnology Information (NCBI) BLAST identification of mutated genes in *Candida albicans* CBS 5982 after NFAP2 or FLC pressure in a microevolution experiment.

### Investigation of cell morphology

Cell morphologies of WT, NFAP2- resistant, and FLC-resistant strains were investigated with scanning electron microscopy to examine whether genomic changes led to morphological alterations (Figure 2A in 15,000× magnification, and Supplementary Figure 3A in 2,000× magnification). Cells of all strains were round-to-oval and secreted extracellular polymeric substance (EPS). EPS secretion, however, was less in NFAP2/6, FLC/5, and FLC/6, compared with the WT. FLC/3 exhibited a wrinkled surface, while the WT and the other resistant strains showed a smooth cell surface (Figure 2A). After NFAP2-treatment (Figure 2B in 15,000× magnification, and Supplementary Figure 3B in 2,000× magnification), WT, NFAP2/4, NFAP2/5, FLC/3, and FLC/5 did not secrete EPS (Figure 2B; Supplementary Figure 3B), and some WT cells collapsed and lost their ovoid shape (Supplementary Figure 3B). Thus, it seems that development of resistance to NFAP2 does not influence cell morphology.

**Figure 2.**
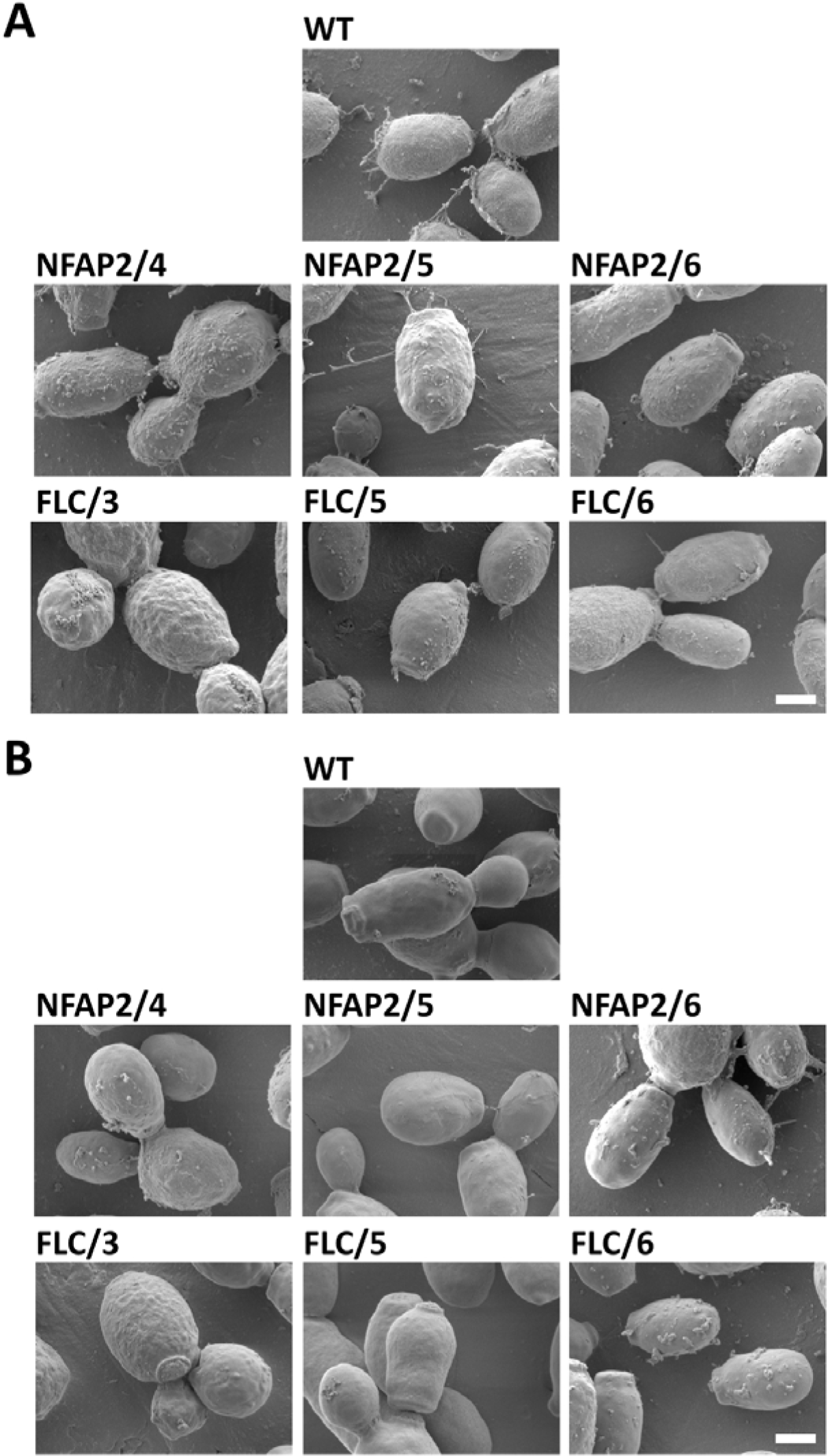
Scanning electron microscopy of the parental wild-type *Candida albicans* CBS 5982 (WT), *Neosartorya* (*Aspergillus*) *fischeri* antifungal protein 2- resistant (NFAP2/4, 5, 6), and fluconazole-resistant (FLC/3, 5, 6) strains (A), and the impact of NFAP2-treatment (1 × minimum inhibitory concentration, 30 min, 30°C, 160 rpm, low-cation medium) on cell morphology (B). 15,000× magnification. The scale bars represent 1 µm.

### Antifungal susceptibility testing to conventional antifungal drugs and antifungal proteins

Development of resistance to an antifungal drug usually results in cross-resistance to other drugs belonging to different antifungal classes (Papp et al., 2020). Antifungal susceptibility tests were performed in LCM to investigate whether the developed resistance influences the susceptibility of *C. albicans* to conventional antifungal agents and ascomycetous anti-*Candida* AFPs. We tested antifungals including the polyene amphotericin B (AMB), triazole FLC, echinocandin micafungin (MFG), allylamine terbinafine (TRB), and NFAP2, as well as AFPs from *Penicillium chrysogenum*, *viz.* PAF, PAFγ^opt^ (Sonderegger et al., 2018), PAFB (Huber et al., 2018), and PAFC (Holzknecht et al., 2020). Susceptibilities of the WT strain to these drugs and AFPs were considered 1 × MIC. Susceptibility tests indicated that the developed resistance did not influence the susceptibilities of NFAP2- and FLC-resistant strains to AMB and MFG and the susceptibility of NFAP2-resistant strains to TBF. Interestingly, the susceptibility of FLC-resistant strains decreased to 2 × MIC of TBF. Regarding FLC, the susceptibilities of NFAP2-resistant strains increased to 0.5 × MIC, and the susceptibilities of FLC-resistant strains decreased to 32 × MIC. PAF and PAFC were ineffective even at the highest investigated concentration (50 µg mL^−1^). The susceptibilities of all resistant strains to PAFγ^opt^ and PAFB were unchanged, except for NFAP2/6, which showed reduced susceptibility to PAFB (2 × MIC). Susceptibility data are summarized in Table 2. Taken together, the susceptibility data indicated that development of resistance to NFAP2 does not influence or increase susceptibility to conventional antifungal drugs, except FLC, and can decrease susceptibility to other AFPs. By contrast, FLC resistance can decrease susceptibility to TRB.

**Table 2.** Minimum inhibitory concentrations (MIC) of conventional antifungal drugs and ascomycetous antifungal proteins (AFPs) in low-cation medium.

| Strain | MIC ($\mu\text{g mL}^{-1}$) | | | | | | | | |
| --- | --- | --- | --- | --- | --- | --- | --- | --- | --- |
| | AMB | FLC | MFG | TBF | NFAP2 | PAF | PAF$\gamma^{\text{opt}}$ | PAFB | PAFC |
| **WT** | 1 | 128 | 0.125 | 64 | 3.125 | >50 | 50 | 25 | >50 |
| **NFAP2/4** | 1 | 64 | 0.125 | 64 | 6.25 | >50 | 50 | 25 | >50 |
| **NFAP2/5** | 1 | 64 | 0.125 | 64 | 6.25 | >50 | 50 | 25 | >50 |
| **NFAP2/6** | 1 | 64 | 0.125 | 64 | 6.25 | >50 | 50 | 50 | >50 |
| **FLC/3** | 1 | 8192 | 0.125 | 128 | 6.25 | >50 | 50 | 25 | >50 |
| **FLC/5** | 1 | 8192 | 0.125 | 128 | 6.25 | >50 | 50 | 25 | >50 |
| **FLC/6** | 1 | 8192 | 0.125 | 128 | 6.25 | >50 | 50 | 25 | >50 |

Susceptibilities of the WT strain to antifungal drugs and AFPs was considered $1 \times \text{MIC}$ . AMB: amphotericin B, FLC: fluconazole, FLC/3, 5, 6: fluconazole-resistant strains, MFG: micafungin, NFAP2: *Neosartorya (Aspergillus) fischeri* antifungal protein 2, NFAP2/4, 5, 6: *Neosartorya (Aspergillus) fischeri* antifungal protein 2-resistant strains, PAF: *Penicillium chrysogenum* antifungal protein, PAF $\gamma^{\text{opt}}$ : γ-core optimized PAF, PAFB: *Penicillium chrysogenum* antifungal protein B, PAFC: *Penicillium chrysogenum* antifungal protein C, TRB: terbinafine, WT: wild-type *Candida albicans* CBS 5982.

### NFAP2 uptake analysis

Studies have suggested that NFAP2 mediates its antifungal activity by disrupting the cell membrane of *C. albicans* (Kovács et al., 2019; Tóth et al., 2018), which requires NFAP2 to bind to the cell. Therefore, we investigated whether reduced NFAP2 binding to the outer cell layers is responsible for a decreased susceptibility to NFAP2 in NFAP2- and FLC-resistant *C. albicans* strains. Therefore, NFAP2 was labeled with the green fluorophore boron-dipyrromethene (BODIPY; Bd-NFAP2) to follow its interaction with the yeast cell. Confocal laser scanning microscopy (CLSM) and fluorescence-activated cell sorting (FACS) analyses were conducted to follow and monitor NFAP2 binding and uptake in *C. albicans*. *C. albicans* cells were co-stained with propidium-iodide (PI) to detect cell death. CLSM analysis showed attachment of Bd-NFAP2 to the outer layers of WT and FLC-resistant *C. albicans* cells in 30 min, followed by accumulation of Bd-NFAP2 in this region or intracellularly, resulting in cell death within 4 h (Supplementary Figure 4). By contrast, attachment of Bd-NFAP2 to the outer cell layers and/or its internalization were not observed or not prominent in NFAP2-resistant cells during this period (Supplementary Figure 4). To quantify *C. albicans* cells that had been treated with BD-NFAP2 and interacted with the antifungal protein, FACS analysis was performed. After 16 h of incubation, FACS results showed a significant decrease in Bd-NFAP2 interaction with each NFAP2-resistant strain (NFAP2/4: 9.26% ± 2.51, NFAP2/5: 18.39% ± 8.66, NFAP2/6: 3.34% ± 2.96) compared with the WT (23.45% ± 0.58) (Table 3). Bd-NFAP2 uptake was significantly increased in FLC/3 (45.51% ± 0.28) and FLC/6 (31.59% 0.69) and significantly decreased in FLC/5 (20.25% ± 1.08). The proportions of dead cells after Bd-NFAP2 uptake are significantly decreased in NFAP2/5 (48.47% ± 4.01), NFAP2/6 (54.43% ± 19.70), FLC/5 (51.43% ± 2.33), and FLC/6 (55.60% ± 1.49), while increased in NFAP2/4 (76.71% ± 9.07), and did not change significantly in FLC/3 (62.49% ± 10.04) compared with the WT (63.03% ± 18.17) (Table 3). These results indicate that development of resistance to NFAP2 and FLC can decrease and increase NFAP2 binding and uptake in *C. albicans*, respectively. FACS data further show that differences in the proportions of dead cell population in Bd-NFAP2-positive cells did not directly correlate with changes in binding/uptake rates between strains.

**Table 3.** *Neosartorya* (*Aspergillus*) *fischeri* antifungal protein 2 (NFAP2) uptake by wild-type, NFAP2- resistant, and FLC-resistant *Candida albicans* strains (%Bd-NFAP2^+^), and the consequent cell death (%PI^+^).

| Strain | Bd-NFAP2+ % | PI+ % |
| --- | --- | --- |
| WT | 23.45 ± 0.58 | 63.03 ± 18.17 |
| NFAP2/4 | 9.26* ± 2.51 | 76.71* ± 9.07 |
| NFAP2/5 | 18.39* ± 8.66 | 48.47* ± 4.01 |
| NFAP2/6 | 3.34* ± 2.96 | 54.43* ± 19.70 |
| FLC/3 | 45.51* ± 0.28 | 62.49 ± 10.04 |
| FLC/5 | 20.25* ± 1.08 | 51.43* ± 2.33 |
| FLC/6 | 31.59* ± 0.69 | 55.60* ± 1.49 |

Bd-NFAP2: BODIPY-labeled, Bd-NFAP2+: Proportion of Bd-NFAP2+ cells in the 30,000 cells detected, FLC/3, 5, 6: fluconazole-resistant strains, NFAP2/4, 5, 6: NFAP2-resistant strains, PI: propidium iodide, PI+: Proportion of PI+ cells in Bd-NFAP2+ detected cells, WT: wild-type *Candida albicans* CBS 5982. \* $p \leq 0.05$ from type II ANOVA followed by Bonferroni's correction as a post hoc test.

### Stress tolerance analyses

Triazole resistance can influence the stress response in *Candida* (Papp et al., 2020); therefore, to study whether development of resistance to NFAP2 influences response to abiotic stress, we conducted spot assays on LCM agar plates. The growth ability of resistant strains was evaluated in the presence of membrane stressors (NaCl and sodium dodecyl sulfate [SDS]) or cell wall stressors (calcofluor white [CFW]), heat treatment or ultraviolet (UV) irradiation compared with the WT. Growth ability of NFAP2/4, NFAP2/6, FLC/5 and FLC/6 was slightly decreased on this medium compared with the WT (Figure 3A). All strains could tolerate the highest applied concentration of membrane stressors (300 mM NaCl and 100 µg mL^−1^ SDS) (Figure 3B) and differences in their growing abilities were not observed. By contrast, NFAP2/4 and NFAP2/6 could not tolerate 10 µg mL^−1^ CFW compared with other strains, whereas FLC/3 had better tolerance (Figure 3B). Below this CFW concentration, tolerance of these two strains was not different from that of the WT (data not shown). Heat treatment at 40°C and 45°C remarkably decreased the growth of NFAP2/4, FLC/3, and FLC/6 compared with the WT (Figure 3C). After heat treatment at 50°C, none of the strains grew on the plates. UV irradiation for 30 s or 60 s decreased or fully inhibited the growth of NFAP2/4, NFAP2/6, and FLC/3 (Figure 3D). Two-minutes-long UV irradiation killed all strains (data not shown). In summary, the development of resistance to NFAP2 can decrease tolerance to cell wall, heat, and UV stresses, whereas development of resistance to FLC can increase tolerance to cell wall stress and decrease tolerance to heat and UV stresses.

**Figure 3.**
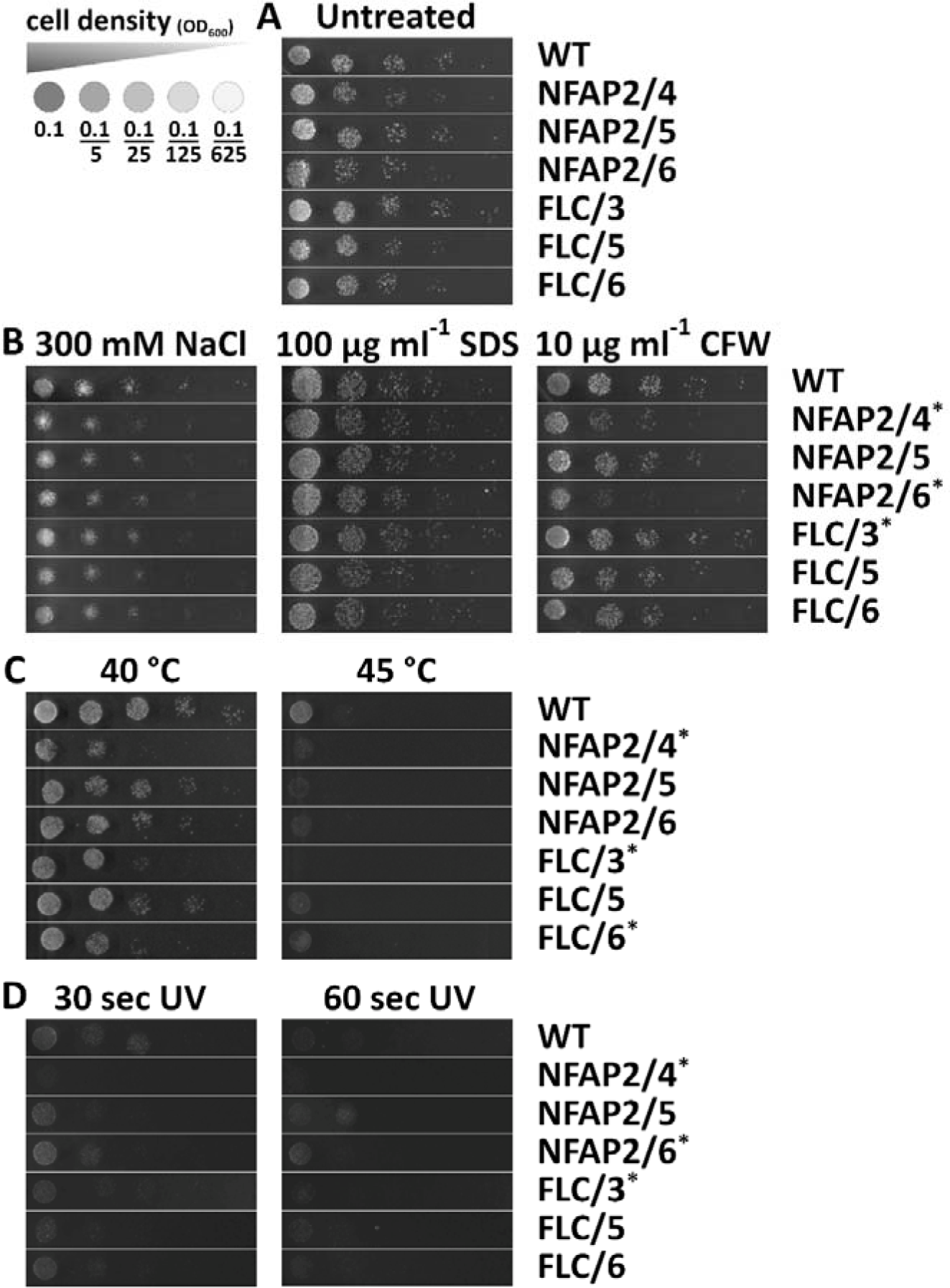
Tolerance of *Candida albicans* strains resistant to *Neosartorya* (*Aspergillus*) *fischeri* antifungal protein 2 (NFAP2/4, 5, 6) and fluconazole (FLC/3, 5, 6) to different abiotic stresses compared with the parental wild-type CBS 5892 (WT) strain. To test growth ability, starting from 0.1 OD_600_ value, five-step 5× serial dilutions of the strains were spotted on low-cation medium (LCM) agar plates without supplementation (A), LCM with plasma membrane (NaCl) or cell wall stressors (SDS: sodium dodecyl sulfate, CFW: calcofluor-white) (B), after 30 min heat treatment at 40°C or 45°C (C), and after ultraviolet (UV) irradiation for 30 or 60 s (D). Gamma settings were adjusted to uniformize background tones for illustrative purposes. *: Change in the stress tolerance compared with WT.

### Metabolic fitness analysis

Development of FLC resistance can reduce the fitness of *C. albicans* in the absence of the drug (Popp et al., 2017); therefore, we investigated whether NFAP2 resistance influences metabolic adaptation to different media compared with the WT. The growth abilities and kinetics of NFAP2- and FLC-resistant strains were monitored in LCM, which was used for the microevolution experiments, malt extract (ME) as rich and Vogel’s as minimal media, and Roswell Park Memorial Institute (RPMI)-1640 as standard clinical susceptibility test medium (Clinical and Laboratory Standards Institute, 2008). Analysis of growth curves indicated that none of the resistant strains had significantly different growing abilities in LCM compared with the WT. The growth of FLC/3 and FLC/5 significantly decreased in ME. In RPMI-1640, each FLC-resistant strain showed significantly weaker growth compared with the WT. In Vogel’s medium, only FLC/5 exhibited reduced growth ability (Figure 4). According to the growth curves, no significant differences were observed in the metabolic adaptation of NFAP2-resistant strains in any medium compared with the WT (Figure 4). These data indicated that resistance to NFAP2 does not have a significant metabolic fitness cost compared with resistance to FLC.

**Figure 4.**
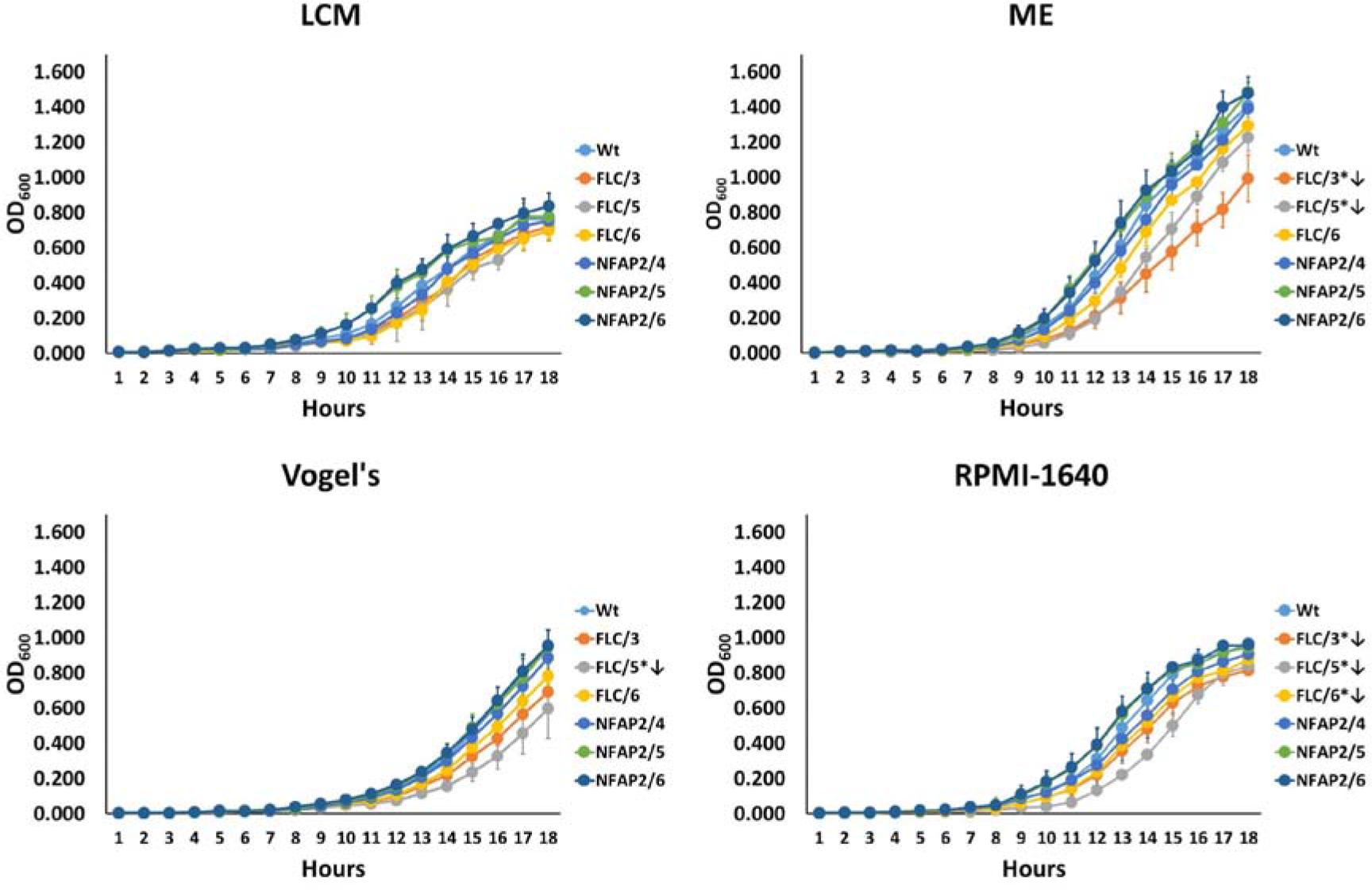
Growth curves of *Candida albicans* strains resistant to *Neosartorya* (*Aspergillus*) *fischeri* antifungal protein 2 (NFAP2/4, 5, 6) and fluconazole (FLC/3, 5, 6) in different media compared with the parental wild-type CBS 5982 (WT) strain (30°C, 160 rpm). LCM: low- cation medium, ME: malt extract medium, Vogel’s: Vogel’s medium, RPMI-1640: Roswell Park Memorial Institute (RPMI)-1640 medium. \**p* ≤ 0.05 from repeated measure ANOVA followed by Tukey–Kramer’s post hoc comparisons. ↓: significantly decreased growth.

### Virulence analysis

Development of resistance to FLC can reduce or increase the virulence of *Candida* species depending on the genomic background of the resistant phenotype (Bohner et al., 2022). Therefore, we investigated the virulence of NFAP2- and FLC**-**resistant strains in a *Galleria mellonella* larval virulence model (Jacobsen, 2014). Survival analysis of larvae infected with resistant *C. albicans* strains indicated that FLC resistance increased (FLC/5 and FLC/6) or decreased (FLC/3) virulence. By contrast, NFAP2 resistance did not influence (NFAP2/4 and NFAP2/5) or decrease (NFAP2/6) virulence (Figure 5). Thus, compared with FLC resistance, NFAP2 resistance does not enhance the virulence of *C. albicans*.

**Figure 5.**
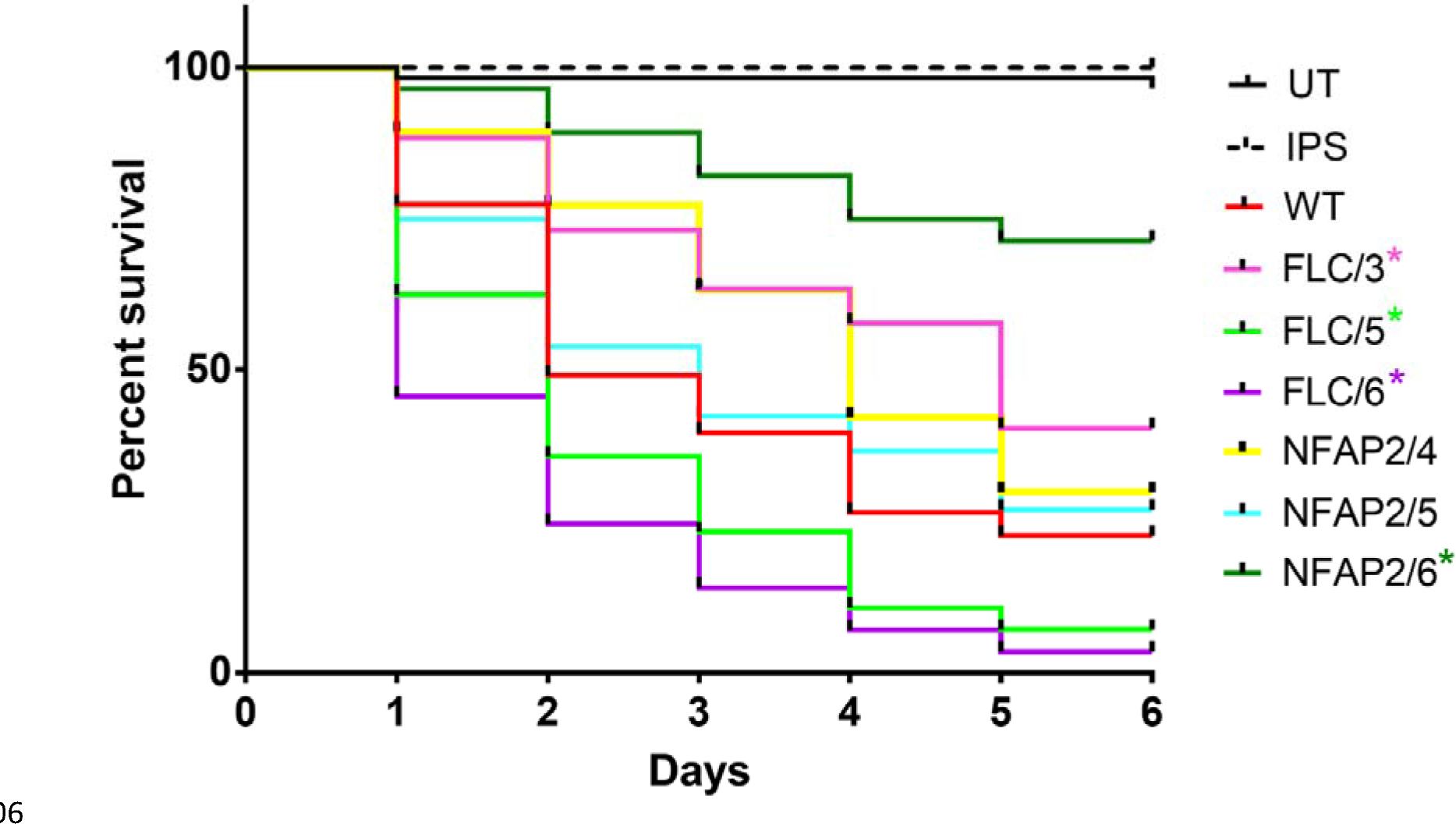
Survival of *Galleria mellonella* larvae after infection with *Candida albicans* strains resistant to *Neosartorya* (*Aspergillus*) *fischeri* antifungal protein 2 (NFAP2/4, 5, 6) and fluconazole (FLC/3, 5, 6) compared with the parental wild-type CBS 5982 (WT) strain. UT: untreated control, IPS: insect physiological saline-treated control. \**p* ≤ 0.05 from both log- rank (Mantel–Cox) and Gehan–Breslow–Wilcoxon tests.

## Discussion

AFPs of filamentous ascomycetes are potent agents for treating topical *Candida* infection (Galgóczy et al., 2019). Several proof-of-concept studies support this hypothesis. These studies showed *in vitro* susceptibilities of various *Candida* spp. to different AFPs, interaction of AFPs with licensed antifungals, potential safe therapeutic applicability of AFPs, and their therapeutic efficacy in mucosal and cutaneous *Candida* infection models (Galgóczy et al., 2019; Holzknecht et al., 2022; Kovács et al., 2019). Despite these promising results, studies on whether *Candida* spp. can develop resistance to fungal AFPs are lacking. These studies are essential to demonstrate the AFPs’ long-term therapeutic applicability in the clinic, where resistance to conventional antifungal drugs is a challenge (Poissy et al., 2022), and new antifungal compounds are needed. Therefore, we examined the potential for developing resistance to one of the most effective AFPs, NFAP2, against the most prevalent human pathogenic yeast, *C. albicans*, compared with generic FLC.

*C. albicans* has remarkable genome plasticity, which helps it overcome the harmful effects of selective environments (Bensasson et al., 2019; Ene et al., 2019). Microevolutionary studies have described the high ability of *C. albicans* to develop strong resistance to FLC, as well as the genomic background of the observed FLC resistance in *C. albicans* (Berkow et al., 2017; Fekete-Forgács et al., 2000; Morschhäuser, 2016). Similar data with AFPs are only available for *Nicotiana alata* defensin NaD1 (McColl et al., 2018) and a human salivary protein histatin 3 (Fitzgerald et al., 2003; Fitzgerald-Hughes et al., 2007) in *Saccharomyces cerevisiae* and *C. albicans*, respectively.

In our experimental setup of microevolution, *C. albicans* could grow even at 32 × MIC of FLC, the highest applied drug concentration, compared with 1 × MIC of NFAP2. This result indicates that *C. albicans* has less ability to develop strong resistance mechanisms to NFAP2, compared with FLC (Figure 1). Consistently, *C. albicans* has been reported to develop resistance to >200 × MIC of FLC (Cowen et al., 2000; Popp et al., 2017; White et al., 1997), while *S. cerevisiae* only adapted to 10 × MIC of NaD1 (McColl et al., 2018), and *C. albicans* became less susceptible to the cell killing effects of histatin 3 (Fitzgerald et al., 2003) in microevolution experiments. Resistance to an antifungal agent can lead to a change in susceptibility to antifungal drugs (Feng et al., 2010; Kelly et al., 1997; McColl et al., 2018; Scott et al., 2023). We observed this in the susceptibility of NFAP2-resistant strains to FLC and FLC-resistant strains to TBF (Table 2). Development of resistance to FLC (Berkow et al., 2017; Morschhäuser, 2016; Popp et al., 2017;) and NaD1 (McColl et al., 2018) is associated with genomic changes that can influence cell morphology, metabolic fitness, stress tolerance, or virulence. We also observed changes in the cell morphology of FLC/3 (Figure 2), metabolic fitness of FLC-resistant strains (Figure 4), stress tolerance of NFAP2/4, NFAP2/6, FLC/3 strains (Figure 3), virulence of NFAP2/6 and all FLC-resistant strains (Figure 5).

The investigation is essential whether the experimental setup of microevolution leads to mutations in functional genes, which could be linked to resistant phenotypes, and results in changes in antifungal susceptibility, metabolic fitness, stress tolerance, and virulence. Therefore, genome analysis of randomly selected FLC- (FLC/3, FLC/5, FLC/6) and NFAP2- resistant (NFAP2/4, NFAP2/5, NFAP2/6) strains were conducted and compared with the genome of the parental WT strain (*C. albicans* CBS 5982) and control lines control lines that had not been treated with the antifungal compounds. Variant analysis showed heterozygous gene mutations in the genome of all resistant strains (Table 1). Considering that *C. albicans* is a diploid yeast, heterozygous mutations can alter cell behavior and modulate drug resistance (Liang et al., 2019).

Studies have reported that genes involved in FLC resistance are typically related to ergosterol biosynthesis (e.g., ERG11, ERG3), efflux pumps (e.g., CDR1, ABC1, MDR1), chaperones (e.g., HSP90), and transcription factors (e.g., UPC2, TAC1) in *C. albicans* (Berkow et al., 2017; Cowen, 2013). Our data obtained with the evolved FLC-resistant strains are not consistent with these reports, since we did not observe mutations in any of the genes mentioned above. The reason of this inconsistency could be the medium and fungal strain differing from that were used in published microevolution experiments to generate FLC resistance in *C. albicans*. However, the mutated genes in our study (CHC1, PTC2, INP51, RIM101) could provide resistance to FLC and could affect susceptibility to TBF, stress tolerance, metabolic fitness, and virulence in *C. albicans*, either directly or indirectly. The respective strain-related phenotypes are discussed below.

FLC/3 contains two SNVs, a nucleotide insertion in CHC1, and two nucleotide deletions in PTC2 (Table 1). Chc1p is a structural subunit of eukaryotic clathrin (Baisa et al., 2013). Clathrin-coated vesicles play a focal role in transfer between cellular compartments; furthermore, clathrin-mediated endocytosis (CME) is the primary mechanism of endocytosis in eukaryotic cells (McMahon et al., 2011). In FLC/3, the nucleotide insertion in CHC1 creates a premature termination codon, which results a partially translated protein without conserved clathrin heavy chain repeat (CLH; smart00299) and clathrin (pfam00637) domains (Supplementary Figure 2). Therefore, the translated protein is most likely dysfunctional and CME is impaired. Rollenhagen et al. (2022) produced a strain of clathrin-deficient *C. albicans* that was resistant to FLC; therefore, we hypothesize that a heterozygous mutation in CHC1 can lead to FLC resistance in FLC/3, because the cellular internalization of FLC is required for its antifungal effect (Rollenhagen et al., 2022). Several publications have indicated that in yeast, clathrin deficiency causes abnormal morphology, slow growth, reduced endocytosis, and weakened cell function, but is not lethal (Goode et al., 2015; Lemmon et al., 1987; Payne et al., 1987; Tan et al., 1993). Our observations regarding FLC/3 can support CHC1 dysfunction, because this strain has a highly wrinkled cell surface (Figure 2A), reduced fitness (Figure 4), heat tolerance (Figure 3C), and virulence (Figure 5). Ptc2p plays a specific role in CO_2_-responsive hyphal elongation in *C. albicans* (Lu et al., 2019). In FLC/3 both nucleotide deletions in PTC2 create a premature termination codon resulting in the loss of 40 C-terminal amino acids of protein (Supplementary Figure 2). The absence of the native C-terminus can potentially compromise the functionality of the protein. CO_2_ is a crucial factor for target location and filamentation in *C. albicans* during host infection when the physiological concentration of CO_2_ can reach several times higher values than that of atmospheric CO_2_ (Bahn et al., 2006; Klengel et al., 2005). Deletion of PTC2 in *C. albicans* caused a defect in CO_2_-induced hyphal elongation at physiological concentrations of CO_2_ (Tan et al., 1993). Mutant Ptc2p, presumed to have diminished function in FLC/3, may contribute to reduced virulence together with the less effective CME (Figure 5).

FLC/5 shows one SNV in a conserved domain region of INP51, which is similar to the catalytic inositol polyphosphate 5-phosphatase (INPP5c) domain of *Saccharomyces cerevisiae* (INPP5c_ScInp51p-like; cd09090) (Supplementary Figure 2). This SNV results in the exchange of a small nonpolar glycine into a hydrophobic valine in the translated protein (Supplementary Figure 2), which can significantly change the physicochemical properties of the protein. Inp51p regulates phosphatidylinositol 4,5-bisphosphate level, maintains cell wall integrity under stress and contact-induced hyphal growth, and may therefore regulate virulence (Badrane et al., 2008; Morales-Johansson et al., 2004). Disruption of Inp51p function in *C. albicans* caused cell wall defects, abnormal distribution of chitin, decreased contact-induced filamentation, and reduced virulence (Badrane et al., 2008). Our observations with FLC/5 contradict these findings, because this strain had a normal cell surface (Figure 2A) and increased virulence in the *G. mellonella* infection model (Figure 5). These results indicate that the glycine-valine change (which increase the overall hydrophobicity) in INP51 does not disrupt the function of Inp51p, but might result in gain-of-function in FLC/5. Considering that FLC causes cell wall stress indirectly through membrane distribution, this gain-of-function mutation may explain the increased FLC tolerance of this strain, because it can compensate the FLC-induced cell wall defect better than WT (Sorgo et al., 2011).

In FLC/6, one SNV is present in a non-conserved domain region of RIM101. This mutation leads to a hydrophobic tyrosine to a hydrophobic cysteine amino acid exchange (Supplementary Figure 2). Rim101p, as a member of the Rim pathway, regulates antifungal drug tolerance proteins, such as Hsp90p and Ipt1p (Garnaud et al., 2018; LaFayette et al., 2010; Prasad et al., 2005; Singh et al., 2009). Deletion of RIM101 increased susceptibility to azoles and echinocandins in *C. albicans* (Cornet et al., 2006). We speculate that the mutation increases the activity of Rim101p, which could explain the FLC resistance of FLC/6.

Studies on the genetic background of tolerance and/or resistance to AFPs are lacking. All studies agree that due to the multimodal mode of action of AFPs (Bleackley et al., 2014; McColl et al., 2018), resistance mechanisms evolve more slowely and require multiple gene mutations compared with conventional antifungal drugs (Katiyar et al., 2009; Sionov et al., 2012). For example, the simultaneous accumulation of mutations in different genes was found to drive NaD1 tolerance in S. cerevisiae because single mutations cannot lead to the observed NaD1-tolerant phenotype (McColl et al., 2018).

The findings of our study show that NFAP2/4 comprises deletion of three nucleotides in an UBIP (GenBank: KHC66060.1) with unknown function and no conserved domains (Supplementary Figure 2). This deletion causes the loss of a polar serine. Considering that this strain showed decreased tolerance to abiotic stress compared with the WT, this gene is assumed to be directly or indirectly involved in stress responses. The genomes of NFAP2/5 and NFAP2/6 contain identical MNVs at the same position in PGA58 (Table 1) with no conserved domain (Supplementary Figure 2). These MNVs result in an amino acid change from a hydrophobic valine to a small nonpolar alanine. Pga58p is involved in cell wall biogenesis, virulence, and biofilm formation in *C. albicans* (Rashid et al., 2022; Silva et al., 2013; Yeater et al., 2007). Because none of the mutations in UBIP and PGA58 in NFAP2- resistant strains are nonsense mutations, the dysfunction of these proteins can be ruled out. All NFAP2-resistant strains showed reduced NFAP2 uptake (Supplementary Figure 4; Table 3), no change in metabolic fitness (Figure 4), and no increase in virulence (Figure 5). Reduced NFAP2 uptake could be caused by changes in the composition and function of the cell wall. Although the NFAP2-resistant strains showed the same susceptibility to NFAP2 in the microdilution assay, the increased cell death of NFAP2/4 after NFAP2 treatment according to FACS analysis (Table 3) and strongly reduced virulence of NFAP2/6 compared with other NFAP2-resistant strains (Figure 5) suggested that *C. albicans* has different ways to develop NFAP2 resistance, which could not be exclusively correlated to the gene mutations observed. These nongenetic processes can be responses to AFP pressure to help overcome its harmful effects, as reported by Fitzgerald-Hughes et al. (2007), who described numerous proteomic changes in histatin 3-resistant derivatives of *C. albicans* compared with the parental strain (Fitzgerald-Hughes et al., 2007). This hypothesis requires further investigation (e.g., transcriptome and proteome analyses) to understand the processes involved in mild NFAP2 resistance.

According to our previous studies, NFAP2 disrupts the plasma membrane at MIC, but does not induce apoptosis in yeast (Tóth et al., 2016). Thus, NFAP2 may have a cell membrane target. However, in this study we observed NFAP2 internalization in *C. albicans* cells treated with sublethal concentrations of NFAP2 (Supplementary Figure 4), indicating that NFAP2 may have an intracellular target. Studying the exact antifungal mechanism of NFAP2 and identification of its molecular target(s) in *C. albicans* can help explain the processes involved in low NFAP2 resistance and the observed phenotypes.

## Conclusion

In this study, we developed resistance to 1 × MIC of NFAP2 and 32 × MIC of FLC in *C. albicans in vitro*, respectively. Genome analysis revealed non-silent mutations in only two genes (UBIP and PGA58) that might be involved in mild NFAP2 resistance. FLC-resistant strains carry mutations in several genes (CHC1, INP51, PTC2, and RIM101), which probably contribute directly or indirectly to resistance and the observed phenotypes. The development of resistance to an antifungal agent leads to changes in susceptibility to other antifungal compounds, fitness, stress tolerance, and virulence. The development of resistance to NFAP2 increased susceptibility to FLC and decreased susceptibility to other AFPs in *C. albicans*. However, the metabolic fitness of NFAP2-resistant *C. albicans* remained unchanged and no increase in virulence was detected compared with FLC-resistant strains. Therefore, *C. albicans* has limited ability to develop strong resistance mechanisms to NFAP2 compared with FLC. NFAP2-based topical anti-*Candida* therapy should be considered, as rapid development of strong resistance may not occur in a short period of time.

## Materials and methods

### Antifungal protein production

Recombinant NFAP2, PAF, PAFγ^opt^, PAFB, and PAFC were produced in a *P. chrysogenum*- based expression system (Sonderegger et al., 2016) and purified, as described (Huber et al., 2018; Holzknecht et al., 2020; Kovács et al., 2019; Sonderegger et al., 2018).

### Microevolution experiment to develop resistance to NFAP2 and FLC

To generate NFAP2- and FLC-resistant strains of *C. albicans* CBS 5982, microevolution experiments were performed in LCM (glucose 5 g L^−1^, yeast extract 0.25 g L^−1^, and peptone 0.125 g L^−1^), supplemented with NFAP2 or FLC (MedChemExpress; Monmouth Junction, NJ, USA). Six independent microevolution experiments were performed in parallel for both. In the adaptation period, mid-log phase *C. albicans* cells (2 × 10^5^ cells mL^−1^) cultured in LCM overnight at 30°C with continuous shaking (160 rpm) were inoculated in a 1:500 volume ratio of LCM supplemented with 0.5 × MIC NFAP2 or FLC and incubated for 24 h (30°C, 160 rpm). The culture was inoculated in a 1:5,000 volume ratio in the same medium and incubated for 24 h two more times. The *Candida* cell cultures were then inoculated in 1:500 volume ratio and the same subculturing steps were applied under the same conditions, but the concentration of NFAP2 or FLC was doubled in the medium (1 × MIC). Serial passage was repeated until the highest antifungal concentration (maximum 32 × MIC) was reached where *Candida* cells could grow. This resulted in *Candida* cell lines that adapted to NFAP2 or FLC. The growth of *Candida* cells was monitored by measuring optical density (OD_600_) of the cultures at the end of the third subculturing step of each antifungal concentration (Genesys 150 UV-Visible Spectrophotometer; Thermo Fischer Scientific, Waltham, MA, USA). In the next evolution period, cultures with the highest concentration of NFAP2 or FLC that *Candida* adapted to were inoculated in a 1:500 volume ratio of LCM without NFAP2 or FLC and incubated for 24 h (30°C, 160 rpm). Then cultures were inoculated in a 1:500 volume ratio of LCM without NFAP2 or FLC and incubated for 24 h. This step was repeated eight more times. Subsequently, 10-fold serial dilutions (10^−1^–10^−6^) of cultures from the last steps were prepared, and 5 µL from the 10^−4^–10^−6^ dilution steps were streaked on 5 mL LCM agar plates (1.5% m/V) (TC Plate 6 Well, Suspension, F; Sarstedt, Nümbrecht, Germany) supplemented with the highest concentrations of NFAP2 or FLC at which *C. albicans* CBS 5982 could grow. The plates were incubated for 48 h at 30°C, then one colony each from the six cell lines was isolated and maintained under selective conditions on LCM agar plates. This phase resulted in six independently evolved NFAP2-, or FLC-resistant *C. albicans* cell lines. The schematic representation of the microevolution experiment is presented in supplementary material (Supplementary Figure 5). As a control, *C. albicans* CBS 5982 was subjected to the same experimental setup, but in the absence of NFAP2 and FLC.

### Whole genome sequencing and data analysis

Genomic DNA was isolated from three mutants, non-selected control, and parental WT strains using Blood & Cell Culture DNA Maxi Kit (QIAGEN, Hilden, Germany) according to the manufacturer’s instructions.

Indexed sequencing libraries were generated from 1 ng genomic DNA using Nextera XT DNA Library Preparation Kit (Illumina, San Diego, CA, USA) following the protocol of the manufacturer. Libraries were validated and quantitated by capillary gel electrophoresis with a 2100 Bioanalyzer instrument (Agilent, Santa Clara, CA, USA), pooled and denatured with 0.1 M NaOH, loaded in MiSeq Reagent Kit v2-300 (Illumina, San Diego, CA, USA) at concentration of 10 pM, and sequenced in a MiSeq DNA sequencing instrument (Illumina, San Diego, CA, USA), generating 2 × 150 bp sequence reads. Primary sequence analysis (base calling, demultiplexing, and fastQ file generation) was performed using Illumina RTA 1.18.54 and GenerateFASTQ 1.1.0.64. The genome of WT *C. albicans* CBS 5982 was assembled with SPAdes 3.13.0 (Nurk et al., 2013) with the help of a reference *C. albicans* SC5314 haploid genome (GCA_000182965.3). Genes were predicted with GlimmerHMM (Majoros et al., 2004).

Variant analysis was performed with CLC Genomics Workbench version 21.0.3 (https://digitalinsights.qiagen.com/). After quality control of reads was complete, the Basic Variant Detection tool was used with default settings. The assembled WT *C. albicans* CBS 5982 genome was used as a reference. Variants with <30% prevalence were filtered out. The variants in WT were removed, and silent mutations were not considered for analysis. Nonsynonymous mutations in exogenic regions are listed in Supplementary File 1. Data availability: The raw reads were uploaded to the European Nucleotide Archive under the bio project No.: PRJEB66720 (https://www.ebi.ac.uk/ena/browser/view/ PRJEB66720).

The acquired gene sequences carrying mutations (Supplementary File 1) were identified using the BLAST tool available on the National Center for Biotechnology Information website (https://blast.ncbi.nlm.nih.gov/Blast.cgi). The resulting gene matches from the *C. albicans* SC5314 genome database with the highest percent identity score above the 90% threshold value were considered identical to the queries (Table 1). Because the reference *C. albicans* SC5314 genome (GCA_000182965.3) used in the assembly of the WT *C. albicans* CBS 5982 genome is available as haploid, in certain cases, variant analysis may recognize native heterozygous loci of the diploid WT genome as mutations. To confirm the presence and heterozygosity of the mutations resulting from genome analyses, the affected gene regions were amplified from the genomic DNA of WT and resistant strains by PCR (SUPPLEMENTARY TABLE 1). The amplicons were subjected to Sanger sequencing (Eurofins Genomics; Ebersberg, Germany) and the obtained sequences were compared with WT sequences through sequence alignment in CLC Genomics Workbench version 21.0.3 (https://digitalinsights.qiagen.com/) software (Sci Ed Software LLC, Westminster, CO, USA). The genes indicated as mutants by WGS were omitted when the amplicon analyses of the WT and the claimed mutant gene regions denied the presence of mutations (highlighted with red in Supplementary File 1). From these gene mutations which were unique to resistant strains and absent in non-selected control lines were considered as a consequence of the NFAP2 or FLC pressure (omitted mutations from non-selected control lines are highlighted in yellow in Supplementary File 1). The conserved domain regions of these mutant genes were identified and subjected to functional analysis by using Conserved Domain Search (https://www.ncbi.nlm.nih.gov/Structure/cdd/wrpsb.cgi; applying default options with “Standard display” result mode) and Conserved Domain Database (https://www.ncbi.nlm.nih.gov/cdd/) (Thanki et al., 2023). Sequence files provided by Eurofins Genomics can be found in Supplementary File 2.

### Cell morphology investigation by scanning electron microscopy

Mid-log phase *C. albicans* cells (4 × 10^6^) were treated with 3.125 µg mL^−1^ NFAP2 in LCM for 30 min at 30°C under continuous shaking at 160 rpm. Untreated cells served as morphology control. Cells were harvested (9,000 × *g* for 5 min), washed two times, and resuspended in phosphate buffered saline (PBS). Then 8 µL samples were spotted on a silicon disk coated with 0.01% (w/v) poly-L-lysine (Merck Millipore, Billerica, MA, USA). Cells were fixed with 2.5% (v/v) glutaraldehyde and 0.05 M cacodylate buffer (pH 7.2) in PBS overnight at 4°C. The disks were washed two times with PBS and dehydrated with a graded ethanol series (30%, 50%, 70%, 80%, and 100% ethanol [v/v] for 24 h each at 4°C). The samples were dried with a Quorum K850 critical-point dryer (Quorum Technologies, Laughton, UK), coated with 12 nm gold, and observed with a JEOL JSM-7100F/LV field emission scanning electron microscope (JEOL Ltd., Tokyo, Japan).

### Antifungal susceptibility testing

According to Tóth et al. (2018) (19), broth microdilution susceptibility tests were performed with conventional antifungal drugs AMB (Santa Cruz Biotechnology, Dallas, TX, USA), FLC (Sigma-Aldrich, St. Louis, MO, USA), MFG (MedChemExpress, Monmouth Junction, NJ, USA), and TRB (MedChemExpress, Monmouth Junction, NJ, USA), as well as AFPs, such as NFAP2, PAF, PAFγ^opt^, PAFB, and PAFC, to determine their MICs in LCM against 2 × 10^4^ cells of WT *C. albicans* CBS5982 and strains resistant to NFAP2 and FLC. Final concentrations were 16–0.0156 μg mL^−1^ for AMB and MFG; 256–0.0156 μg mL^−1^ for TRB; 8192–0.0625 μg mL^−1^ for FLC; and 50–1.562 μg mL^−1^ for NFAP2, PAF, PAFγ^opt^, PAFB, and PAFC in twofold dilutions. The cultures were resuspended by pipetting in microtiter plates (TC Plate 96 Well, Suspension, F; Sarstedt, Nümbrecht, Germany) and incubated at 30°C for 48 h without shaking. OD_600_ values were measured with a microplate reader (Thermo Labsystems 354 Multiskan Ascent Microplate Reader; Thermo Fischer Scientific, Waltham, MA, USA) after suspending the cells in each well. MIC was defined as the lowest concentration of an antifungal compound that reduces fungal growth to 10% compared with the untreated control, which was set to 100%. Antifungal susceptibility tests were repeated at least two times, including two technical replicates.

### NFAP2 localization studies

For NFAP2 localization in *C. albicans*, NFAP2 was labeled with the green fluorophore BODIPY FL-EDA (Bd; Invitrogen, Waltham, MA, USA) according to Sonderegger et al. (2017) (Sonderegger et al., 2017). Mid-log phase *C. albicans* cells (4 × 10^6^) were treated with 2.5 µg mL^−1^ Bd-NFAPP2 in LCM for 30, 60, 120, and 180 min at 30°C with continuous shaking at 160 rpm, washed with PBS, and incubated with 5 µg mL^−1^ propidium iodide (PI; Sigma-Aldrich, St. Louis, MO, USA) for 10 min at room temperature in the dark, washed two times with PBS, and resuspended in PBS. The samples were examined with a confocal laser (488 nm) scanning microscope (Olympus Fluoview FV 1000; Olympus, Shinjuku, Japan). Excitation and emission wavelengths were 504 and 512 nm for NFAP2-Bd and 535 and 617 nm for PI, respectively. Sequential scanning was used to avoid crosstalk between the fluorescent dyes.

### NAFP2-uptake analysis

NFAP2 uptake by *C. albicans* strains was analyzed by FACS. *C. albicans* cells (4 × 10^6^) were treated with 3.125 µg mL^−1^ Bd-NFAP2 in LCM (30°C, 16 h, 160 rpm) and costained with PI (as described for NFAP2 localization). Bd-NFAP2- and PI-positive cells were detected by a FlowSight imaging flow cytometer equipped with lasers at 405 (violet), 488 (blue), and 642 nm (red) (Amins, Merck Millipore, Billerica, MA, USA). Calibration controls were used to avoid overexposure of the positive events in specific fluorescent channels, which cause false positive staining signals in other fluorsecent channels. Cells treated with 70% (v/v) ethanol (10 min at room temperature, 160 rpm) were used as positive PI staining and calibration controls, and cells treated with 2 µg mL^−1^ fluorescein diacetate (Thermo Fischer Scientific, Waltham, MA, USA) (20 min at room temperature, 160 rpm) were used as Bd-NFAP2 calibration controls. 10,000 cells per run were detected. Bd-NFAP2 was detected at 488 nm and PI at 642 nm, with excitation lasers and emission in channel 2 window. Gating was adjusted to reach at least 96% of the untreated cells and debris was excluded during data acquisition. Data analysis was performed with Image Data Exploration and Analysis software (IDEAS; Amins, Millipore, Billerica, MA, USA). The FACS experiments were repeated three times.

### Stress tolerance analyses

For stress tolerance assays, *C. albicans* strains were grown overnight in LCM (30°C, 160 rpm) to reach the mid-log phase, the OD_600_ of the cultures was set at 0.1, and fivefold dilution series were prepared in the same medium. Five microliters from the dilution series were spotted on the surface of LCM agar plates and dried before incubation at 30°C for 24 h. After the incubation period, the plates were photographed (Versa Doc Imaging System 4000 MP; Bio-Rad, Hercules, CA, USA). To investigate tolerance to NaCl, SDS, and CFW, agar plates were supplemented with serial dilutions of CFW (1, 2.5, 5, and 10 µg mL^−1^), SDS (12.5, 25, 50, and 100 µg mL^−1^), or NaCl (50, 100, 200, and 300 mM). To investigate tolerance to UV irradiation, the plates were irradiated with UV light for 0, 0.5, 1, 2, and 5 min with the germicidal lamp of a laminar flow box (Herasafe; Thermo Electron Corporation, Waltham, MA, USA). To investigate tolerance to heat stress, the fivefold dilution series of *C. albicans* strains were heat-treated at 30°C, 37°C, 40°C, and 45°C for 30 min. Two technical replicates were prepared for all stress tolerance analyses and tests were repeated at least two times.

### Metabolic fitness assays

Metabolic adaptation and growth of WT CBS 5982 and NFAP2- and FLC-resistant strains of *C. albicans* were monitored in LCM, Vogel’s medium (10 g L^−1^ D-glucose, 2.5 g L^−1^ Na_3_- citrate, 5 g L^−1^ anhydrous KH_2_PO_4_, 2 g L^−1^ anhydrous NH_4_NO_3_, 0.2 g L^−1^ MgSO_4_L7H_2_O, 0.1 g L^−1^ CaC1_2_L2H_2_O, 5.26 mg L^−1^ monohydrous citric acid, 5.26 mg L^−1^ ZnSO_4_L7H_2_O, 1.05 mg L^−1^ Fe(NH_4_)_2_(SO_4_)L6H_2_O, 0.26 mg L^−1^ CuSO_4_L5H_2_O, 0.05 mg L^−1^ MnSO_4_L4H_2_O, 0.05 mg L^−1^ H_3_BO_3_, 0.05 mg L^−1^ Na_2_MoO_4_L2H_2_O, and 0.05 mg L^−1^ biotin), ME (Sigma-Aldrich, St. Louis, MO, USA), and RPMI-1640 (Gibco - Thermo Fischer Scientific, Waltham, MA, USA) media and compared with that of WT. Next, 200 µL of mid-log phase *C. albicans* cells (OD_600_ = 0.1) were inoculated in 20 mL of the respective media and incubated for 18 h at 30°C and 160 rpm. OD_600_ values of the cultures were measured every hour with a Genesys 150 UV-Visible Spectrophotometer (Thermo Fischer Scientific, Waltham, MA, USA). The metabolic fitness assays were repeated three times.

### Virulence assay

A *G. mellonella in vivo* infection model was used to investigate the virulence of the generated NFAP2- and FLC-resistant strains compared with the parental WT CBS 5982 strain. Twenty *G. mellonella* larvae (TruLarv; BioSystems Technology, Exeter, United Kingdom) were infected with 20 µL of 5 × 10^7^ *C. albicans* cells mL^−1^ suspended in insect physiological saline (IPS: 50 mM NaCl, 5 mM KCl, 10 mM EDTA, and 30 mM sodium citrate in 0.1 M Tris-HCl [pH 6.9]) by intrahemocoelic injection (29-gauge insulin needles; BD Micro-Fine, Franklin Lakes, NJ, USA) through the last proleg. IPS-treated larvae served as the uninfected controls, whereas larvae without interventions served as untreated controls. The larvae were incubated at 37°C, and survival was monitored every 24 h for 6 d. The virulence assay was repeated three times.

### Statistical analyses

For FACS analysis, a generalized linear regression model with binomial distribution was applied to the data due to the categorical nature of the response variable (glm() function). Models were evaluated using type II ANOVA. Pairwise comparison with Bonferroni’s correction was used for post hoc analysis in cases of significant effects. During pairwise comparison, estimated marginal means were tested against the dataset of the WT; *p* values ≤ 0.05 were considered significant (n = 30,000 per strain). To assess the fitness curves of the different strains, repeated measure ANOVA was used (fitrm() and ranova() functions), and a different repeated measure model was fitted for each substrate. In the case of significant models, the Tukey–Kramer post hoc test was used for comparison; *p* values ≤ 0.05 were considered significant. MATLAB 2022b (MathWorks, Portola Valley, CA, USA) and R (R Core Team (2022) R: A Language and Environment for Statistical Computing, R Foundation for Statistical Computing, Vienna) software were used to evaluate FACS and fitness test data. To compare survival curves in *G. mellonella* larval infection model experiments, log-rank (Mantel–Cox) and Gehan–Breslow–Wilcoxon tests were used in GraphPad Prism 7.00 (GraphPad Software, Boston, MA, USA). Survival was considered significant if *p* ≤ 0.05 in both tests, n = 60 per strain.

## Supplemental material

Supplemental material is available online only.

## Acknowledgments

The present work of L.G. was financed by the Hungarian National Research, Development and Innovation Office - NKFIH, FK 134343 and K 146131 projects. The research was funded in part by the Austrian Science Fund FWF (I3132-B21) to F.M. University of Szeged Open Access Fund, Grant ID: 7014.

## Supplementary materials

**Supplementary Table 1.**
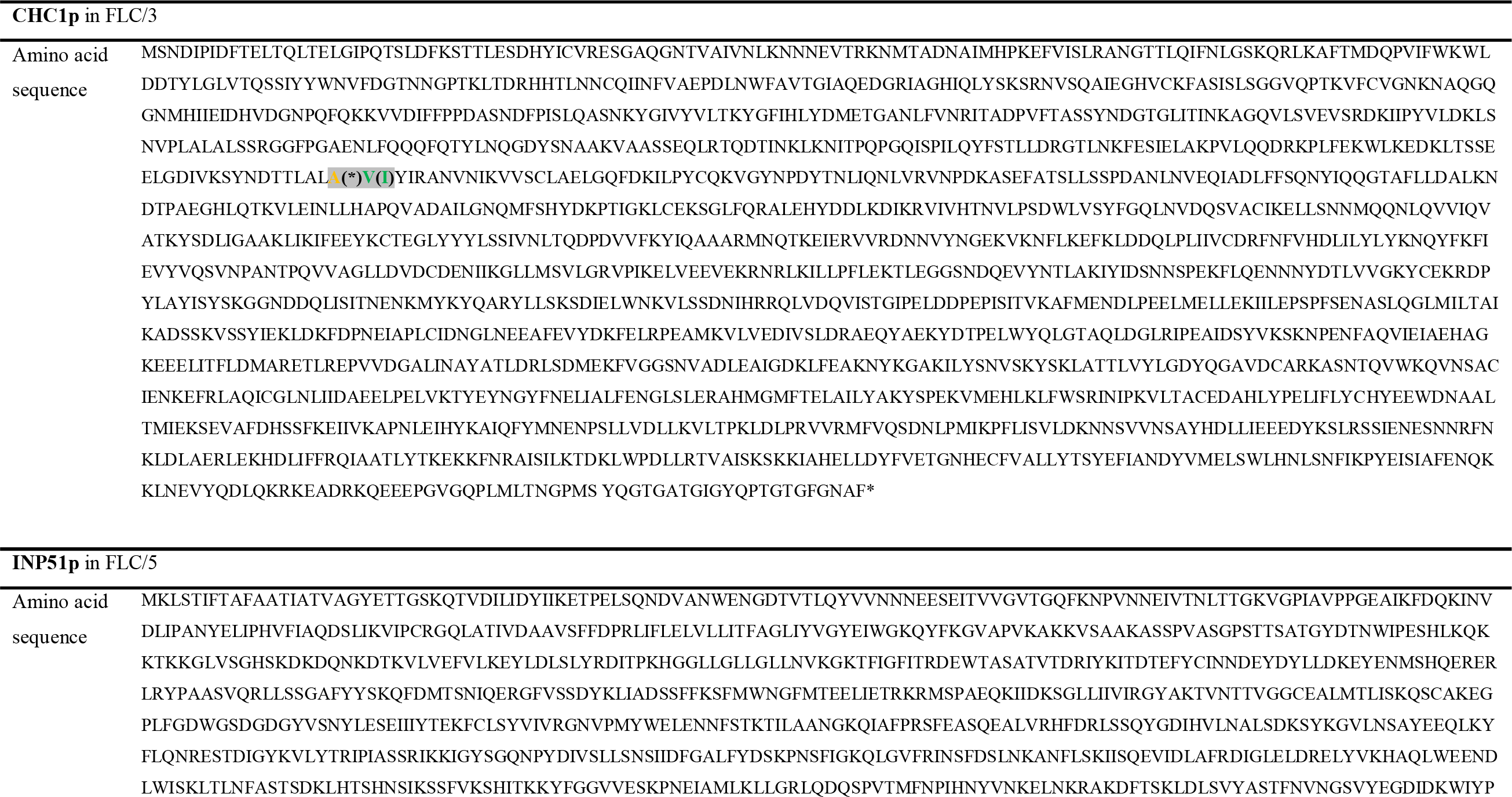
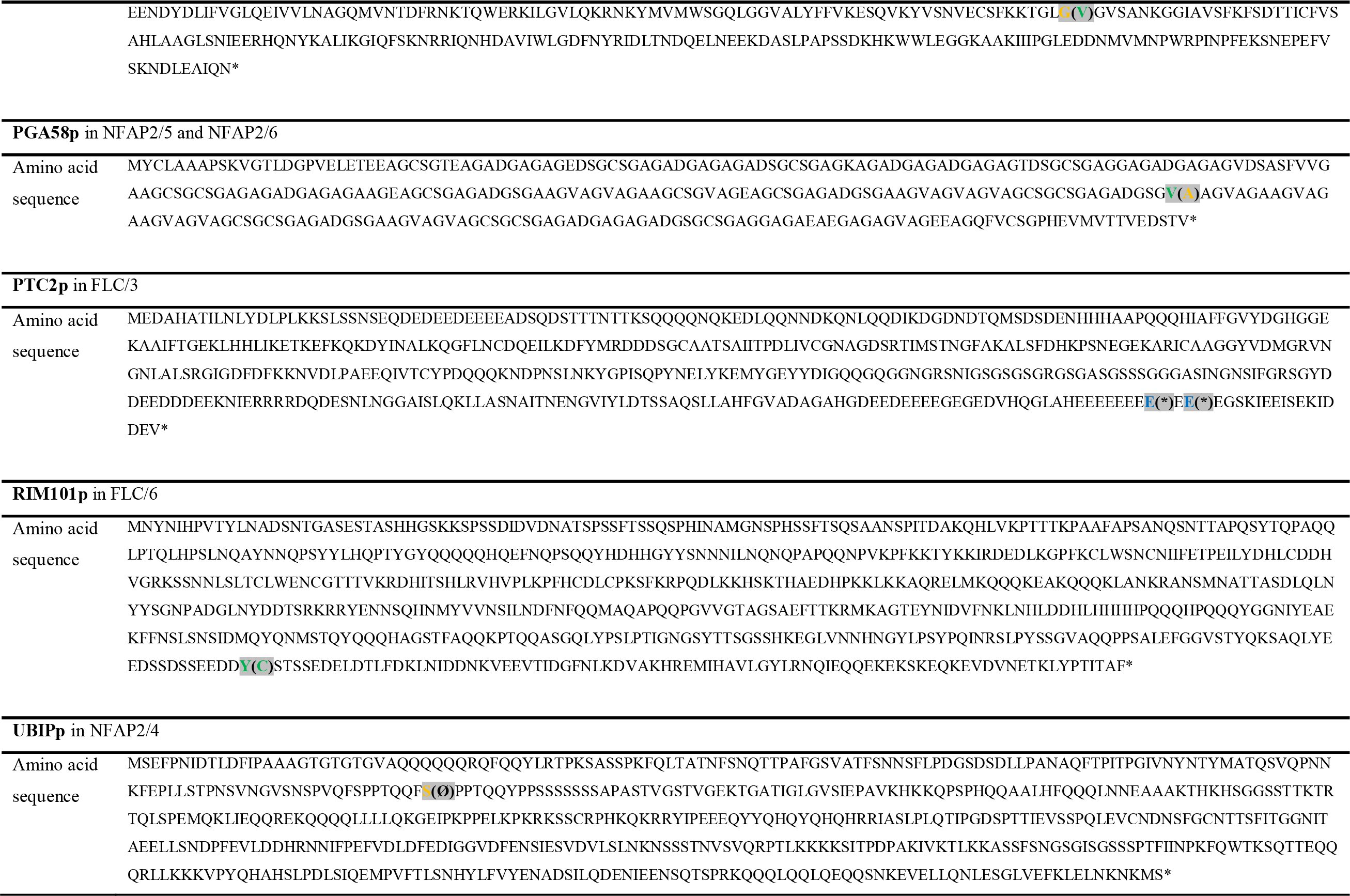
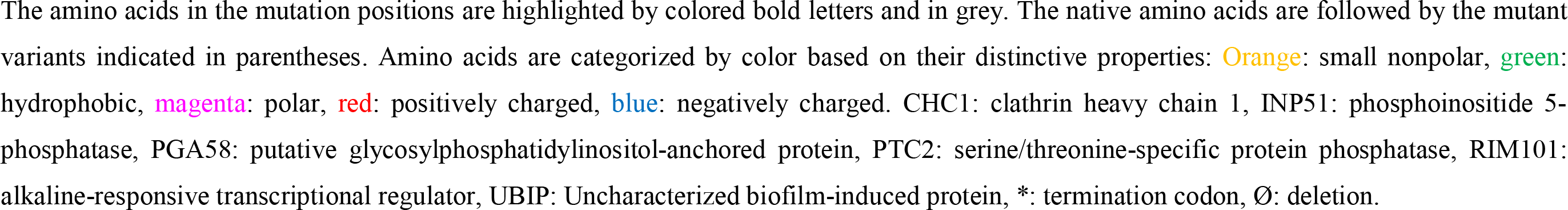
Positions of mutations in the amino acid sequences of the affected proteins in *Neosartorya* (*Aspergillus*) *fischeri* antifungal protein 2 (NFAP2/4, 5, 6) and fluconazole (FLC/3, 5, 6) resistant strains.

**Supplementary Table 2.**
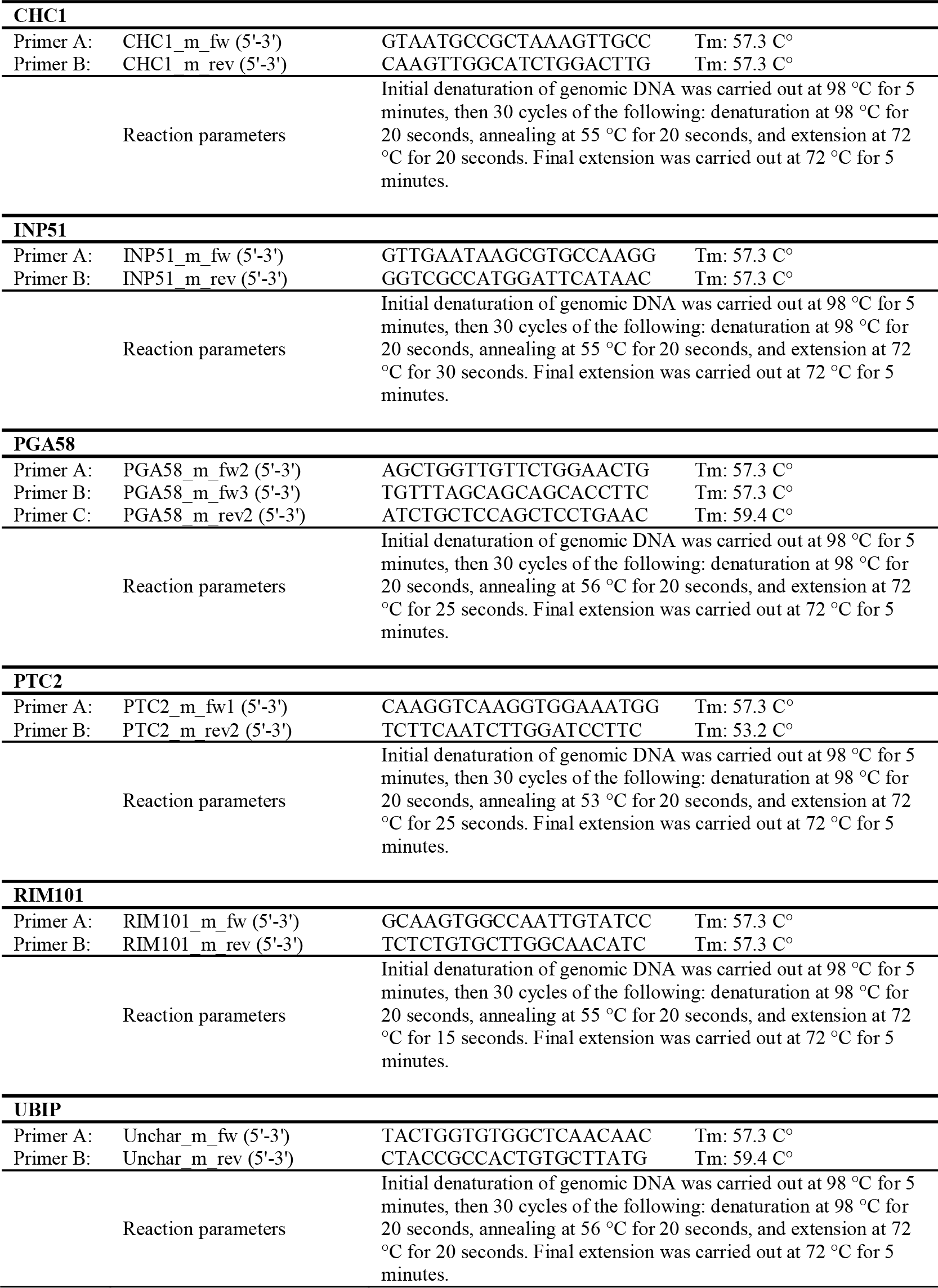
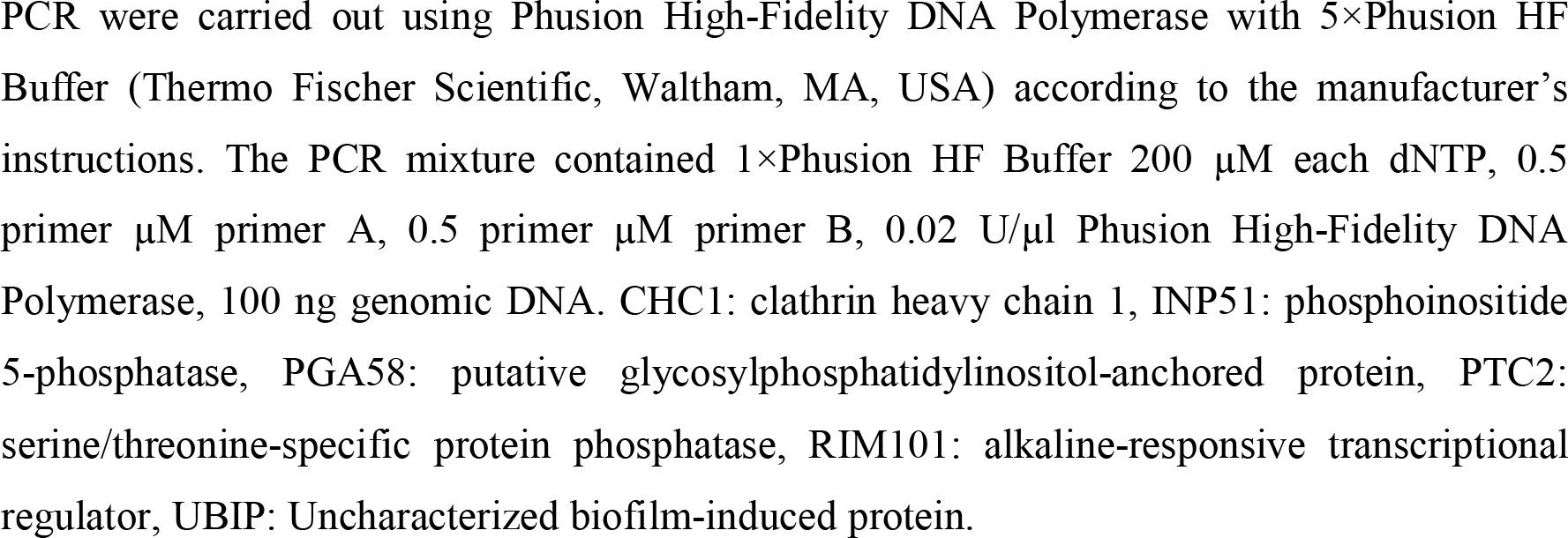
Oligonucleotides and reaction parameters of PCR applied for amplification of the mutated gene regions in the consequence of NFAP2 or FLC pressure in a microevolution experiment.

**Supplementary Figure 1.**
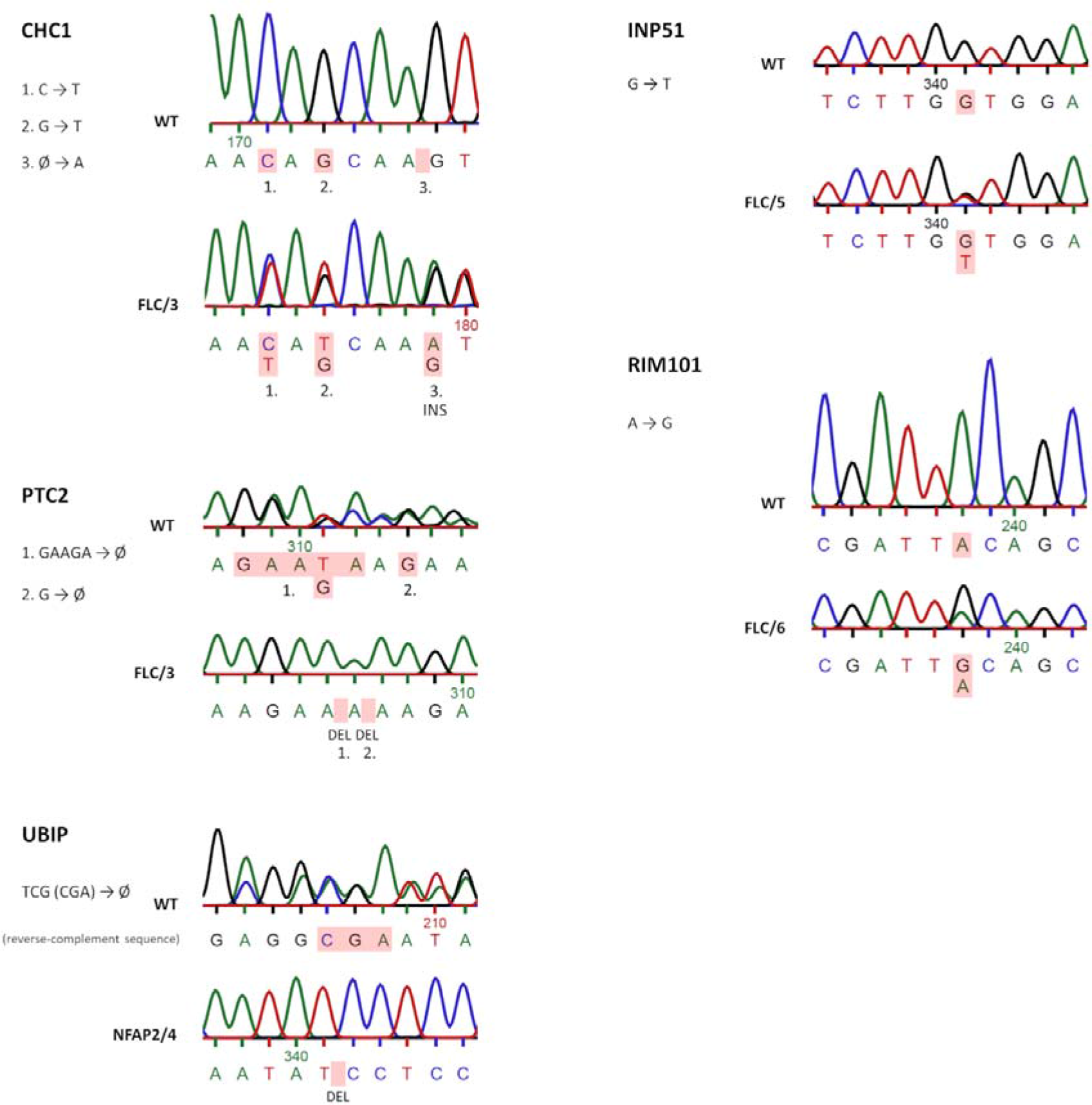
Sequenograms of the mutated gene regions in the consequence of *Neosartorya* (*Aspergillus*) *fischeri* NRRL 181 antifungal protein 2 (NFAP2) or fluconazole (FLC) pressure in a microevolution experiment in comparison with the related regions of the wild-type (WT) *Candida albicans* CBS 5982 genes. Base changes are highlighted in red. Double peaks in the mutated region indicate heterozygous mutations. CHC1: clathrin heavy chain 1, DEL: deletion, FLC/3, 5, 6: FLC-resistant strains, INP51: phosphoinositide 5- phosphatase, INS: insertion, NFAP2/4, 5, 6: NFAP2-resistant strains, PTC2: serine/threonine- specific protein phosphatase, RIM101: alkaline-responsive transcriptional regulator, UBIP: Uncharacterized biofilm-induced protein. The full length sequenograms files can be found in Supplementary File 2.

**Supplementary Figure 2.**
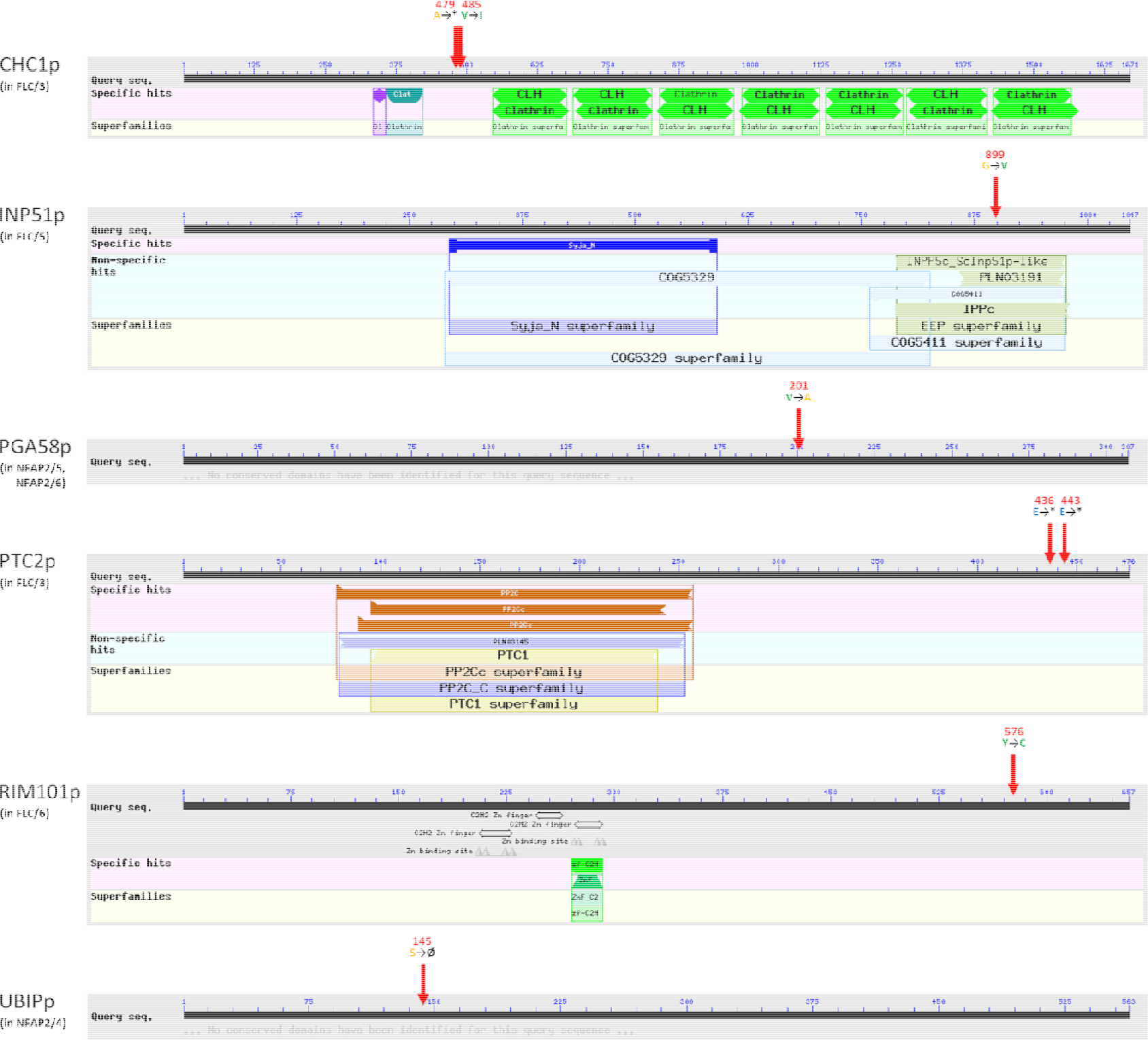
Locations of detected *Candida albicans* CBS 5982 gene mutations at amino acid level as consequences of *Neosartorya* (*Aspergillus*) *fischeri* NRRL 181 antifungal protein 2 (NFAP2) or fluconazole (FLC) pressure. Conserved domains are indicated below query seq. The specific positions (amino acid number) of respective mutations in the sequence are indicated with red arrows. The amino acid alterations are indicated with colored letters. Amino acids are categorized by color based on their distinctive properties: Orange small nonpolar, green: hydrophobic, magenta: polar, red: positively charged, blue: negatively charged. CHC1: clathrin heavy chain 1, INP51: phosphoinositide 5- phosphatase, PGA58: putative glycosylphosphatidylinositol-anchored protein, PTC2: serine/threonine-specific protein phosphatase, RIM101: alkaline-responsive transcriptional regulator, UBIP: Uncharacterized biofilm-induced protein, *: termination codon, Ø: deletion. Graphical summaries were generated by Conserved Domain Search (https://www.ncbi.nlm.nih.gov/Structure/cdd/wrpsb.cgi; default options with “Standard display” result mode) using Conserved Domain Database (https://www.ncbi.nlm.nih.gov/cdd/).

**Supplementary Figure 3.**
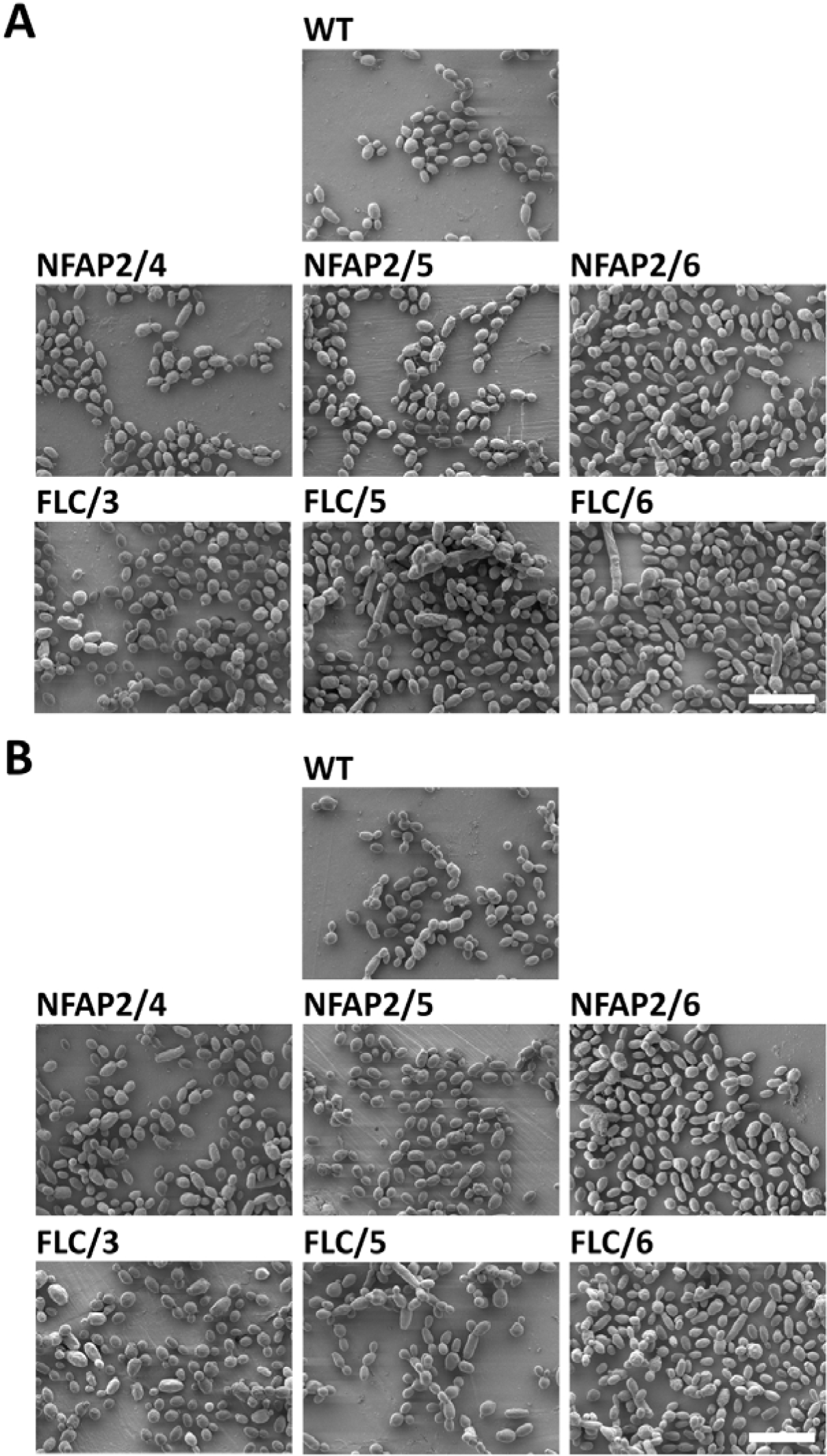
Scanning electron microscopy of the parental wild-type *Candida albicans* CBS 5982 (WT), *Neosartorya* (*Aspergillus*) *fischeri* antifungal protein 2- resistant (NFAP2/4, 5, 6), and fluconazole-resistant (FLC/3, 5, 6) strains (A), and the impact of NFAP2-treatment (1 × minimum inhibitory concentration, 30 min, 30°C, 160 rpm, low-cation medium) on cell morphology (B). 2,000× magnification. Scale bars represent 10 µm.

**Supplementary Figure 4.**
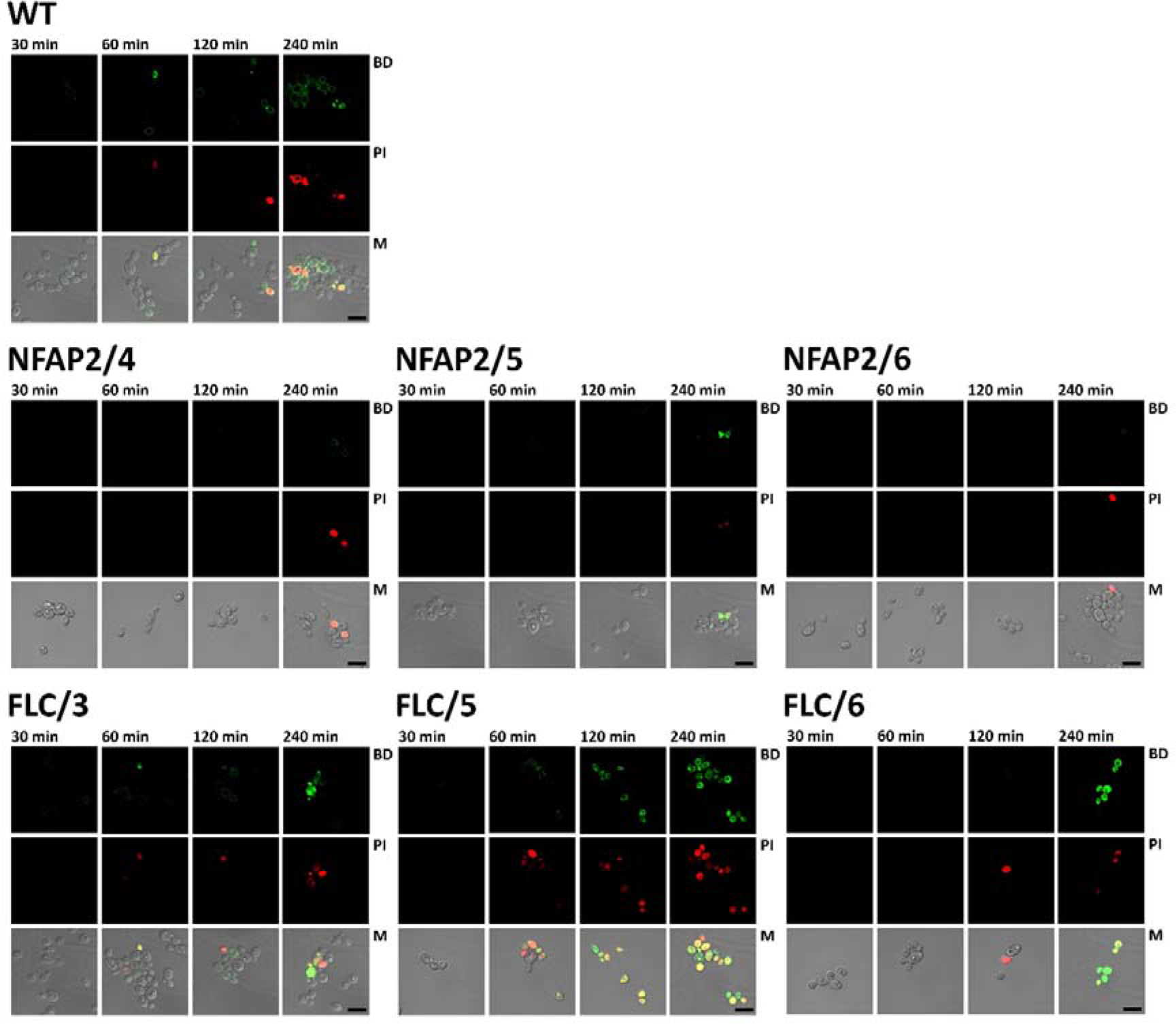
Confocal laser scanning microscopy of the parental wild-type CBS 5982 (WT), NFAP2-resistant (NFAP2/4, 5, 6), and fluconazole-resistant (FLC/3, 5, 6) *Candida albicans* CBS 5982 strains (4 × 10^6^ cells) after treatment with the green fluorophore BODIPY-labelled NFAP2 (Bd-NFAP2, 3.125 µg ml^−1^) for 30, 60, 120, and 240 minutes (30°C, 160 rpm, low cationic medium), and co-stained with propidium-iodide (PI) for 10 minutes. Sequential scanning was done for Bd-NFAP2 (BD, green) and PI (PI, red) at 488 and 543 nm, respectively. M: merged BP, PI, and bright field pictures. Scale bars represent 10 µm. The gamma settings of the merged images have been adjusted to uniformize the tones of the backgrounds for illustrative purposes.

**Supplementary Figure 5.**
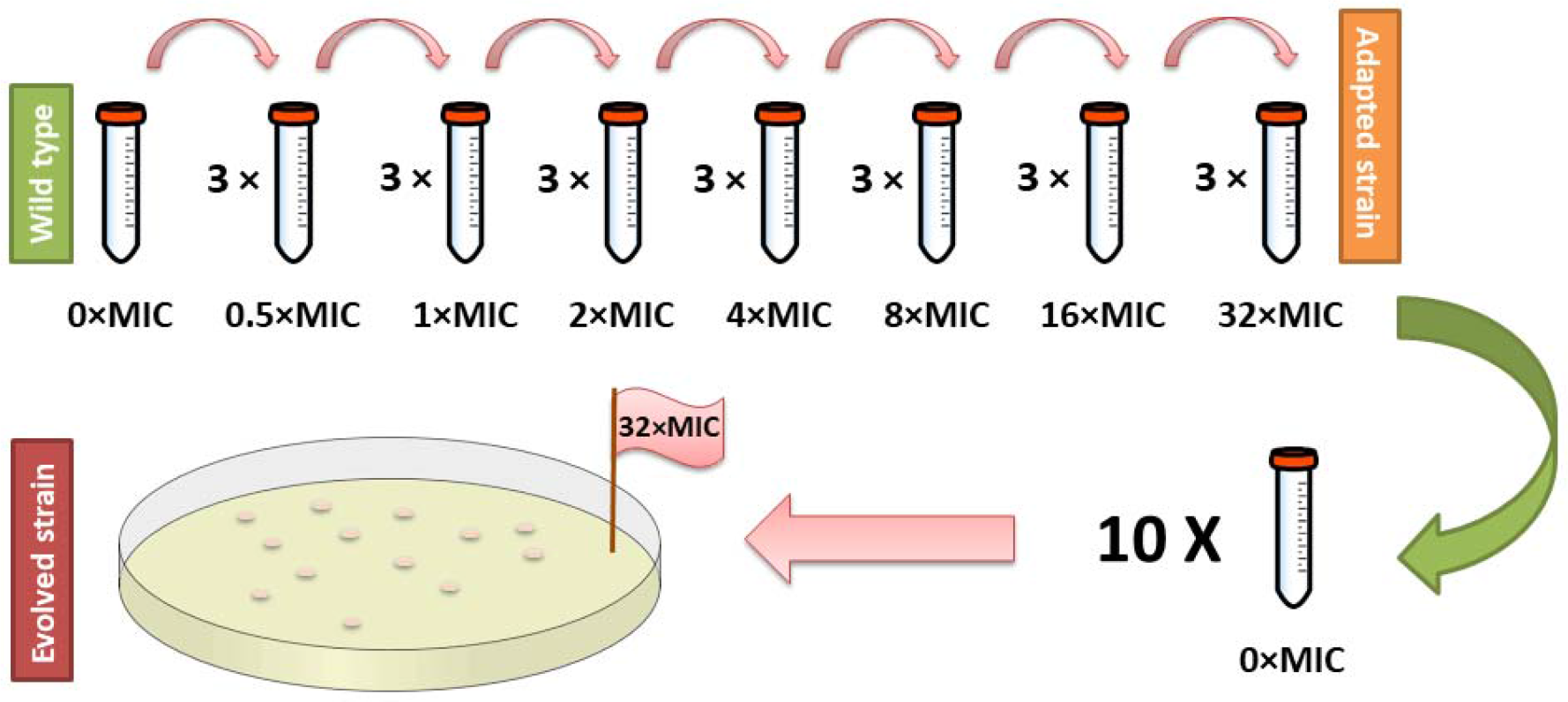
Schematic representation of microevolution experiment performed in low cationic medium to generate NFAP2- or fluconazole-resistant *Candida albicans* strains.

## References

1. Alexander BD, Johnson MD, Pfeiffer CD, Jiménez-Ortigosa C, Catania J, Booker R, Castanheira M, Messer SA, Perlin DS, Pfaller MA. 2013. Increasing echinocandin resistance in *Candida glabrata*: clinical failure correlates with presence of FKS mutations and elevated minimum inhibitory concentrations. Clin Infect Dis 56:1724–1732. DOI: 10.1093/cid/cit136.

2. Augostine CR, Avery SV. 2022. Discovery of natural products with antifungal potential through combinatorial synergy. Front Microbiol 13:866840. DOI: 10.3389/fmicb.2022.866840.

3. Badrane H, Nguyen MH, Cheng S, Kumar V, Derendorf H, Iczkowski KA, Clancy CJ. 2008. The *Candida albicans* phosphatase Inp51p interacts with the EH domain protein Irs4p, regulates phosphatidylinositol-4, 5-bisphosphate levels and influences hyphal formation, the cell integrity pathway and virulence. Microbiology 154:3296–3308. DOI: 10.1099/mic.0.2008/018002-0.

4. Bahn YS, Mühlschlegel FA. 2006. CO_2_ sensing in fungi and beyond. Curr Opin Microbiol 9:572–578. DOI: 10.1016/j.mib.2006.09.003.

5. Baisa GA, Mayers JR, Bednarek SY. 2013. Budding and braking news about clathrin-mediated endocytosis. Curr Opin Plant Biol 16:718–725. DOI: 10.1016/j.pbi.2013.09.005.

6. Bensasson D, Dicks J, Ludwig JM, Bond CJ, Elliston A, Roberts IN, James SA. 2019. Diverse lineages of *Candida albicans* live on old oaks. Genetics 211:277–288. DOI: 10.1534/genetics.118.301482.

7. Berkow EL, Lockhart SR. 2017. Fluconazole resistance in *Candida* species: a current perspective. Infect Drug Resist. 10:237–245. DOI: 10.2147/IDR.S118892.

8. Bleackley MR, Wiltshire JL, Perrine-Walker F, Vasa S, Burns RL, van der Weerden NL, Anderson MA. 2014. Agp2p, the plasma membrane transregulator of polyamine uptake, regulates the antifungal activities of the plant defensin NaD1 and other cationic peptides. Antimicrob Agents Chemother 58:2688–2698. DOI: 10.1128/AAC.02087-13.

9. Bohner F, Papp C, Gácser A. 2022. The effect of antifungal resistance development on the virulence of *Candida* species. FEMS Yeast Res 22:foac019. DOI: 10.1093/femsyr/foac019.

10. Bongomin F, Gago S, Oladele RO, Denning DW. 2017. Global and multi-national prevalence of fungal diseases-estimate precision. J Fungi 3:57. DOI: 10.3390/jof3040057.

11. Clinical and Laboratory Standards Institute. Reference Method for Broth Dilution Antifungal Susceptibility Testing of Yeasts. Approved Standard, 3rd ed.; M27-A3; CLSI: Wayne, PA, USA, 2008

12. Cornet M, Gaillardin C, Richard ML. 2006. Deletions of the endocytic components VPS28 and VPS32 in *Candida albicans* lead to echinocandin and azole hypersensitivity. Antimicrob Agents Chemother 50:3492–3495. DOI: 10.1128/AAC.00391-06.

13. Cowen LE, Sanglard D, Calabrese D, Sirjusingh C, Anderson JB, Kohn LM. 2000. Evolution of drug resistance in experimental populations of *Candida albicans*. J Bacteriol 182:1515– 1522. DOI: 10.1128/JB.182.6.1515-1522.2000.

14. Cowen LE. 2013. The fungal Achilles’ heel: targeting Hsp90 to cripple fungal pathogens. Current Opinion in Microbiology 16;4: 377–384. DOI: 10.1016/j.mib.2013.03.005.

15. Ene IV, Bennett RJ, Anderson MZ. 2019. Mechanisms of genome evolution in *Candida albicans*. Curr Opin Microbiol. 52:47–54. DOI: 10.1016/j.mib.2019.05.001.

16. Fekete-Forgács K, Gyüre L, Lenkey B. 2000. Changes of virulence factors accompanying the phenomenon of induced fluconazole resistance in *Candida albicans*. Mycoses. 43:273–279. DOI: 10.1046/j.1439-0507.2000.00587.x.

17. Feng LJ, Wan Z, Wang XH, Li RY, Liu W. 2010. Relationship between antifungal resistance of fluconazole resistant *Candida albicans* and mutations in ERG11 gene. Chin Med J (Engl) 123:544–548. DOI: 10.3760/cma.j.issn.0366-6999.2010.05.007.

18. Fernández de Ullivarri M, Arbulu S, Garcia-Gutierrez E, Cotter PD. 2020. Antifungal peptides as therapeutic agents. Front Cell Infect Microbiol 10:105. DOI: 10.3389/fcimb.2020.00105.

19. Fitzgerald DH, Coleman DC, O’Connell BC. 2003. Binding, internalisation and degradation of histatin 3 in histatin-resistant derivatives of *Candida albicans*. FEMS Microbiol Lett 220:247–253. DOI: 10.1016/S0378-1097(03)00121-6

20. Fitzgerald-Hughes DH, Coleman DC, O’Connell BC. 2007. Differentially expressed proteins in derivatives of *Candida albicans* displaying a stable histatin 3-resistant phenotype. Antimicrob Agents Chemother 51:2793–800. DOI: 10.1128/AAC.00094-07.

21. Friedman DZP, Schwartz IS. 2019. Emerging fungal infections: New patients, new patterns, and new pathogens. J Fungi (Basel). 5:67. DOI: 10.3390/jof5030067.

22. Galgóczy L, Yap A, Marx F. 2019. Cysteine-rich antifungal proteins from filamentous fungi are promising bioactive natural compounds in anti-*Candida* therapy. Isr J Chem 59:360–370. DOI: 10.1002/ijch.201800168.

23. Garnaud C, García-Oliver E, Wang Y, Maubon D, Bailly S, Despinasse Q, Champleboux M, Govin J, Cornet M. 2018. The Rim pathway mediates antifungal tolerance in *Candida albicans* through newly identified Rim101 transcriptional targets, including Hsp90 and Ipt1. Antimicrob Agents Chemother 62:e01785–17. DOI: 10.1128/AAC.01785-17.

24. Goode BL, Eskin JA, Wendland B. 2015. Actin and endocytosis in budding yeast. Genetics 199:315–358. DOI: 10.1534/genetics.112.145540.

25. Govindarajan A, Bistas KG, Ingold CJ, Aboeed A. 2023. Fluconazole. In: StatPearls [Internet]. Treasure Island (FL): StatPearls Publishing.

26. Holzknecht J, Dubrac S, Hedtrich S, Galgóczy L, Marx F. 2022. Small, cationic antifungal proteins from filamentous fungi inhibit *Candida albicans* growth in 3D skin infection models. Microbiol Spectr 10:e0029922. DOI: 10.1128/spectrum.00299-22.

27. Holzknecht J, Kühbacher A, Papp C, Farkas A, Váradi G, Marcos JF, Manzanares P, Tóth GK, Galgóczy L, Marx F. 2020. The *Penicillium chrysogenum* Q176 antimicrobial protein PAFC effectively inhibits the growth of the opportunistic human pathogen *Candida albicans*. J Fungi (Basel) 6:141. DOI: 10.3390/jof6030141.

28. Hossain CM, Ryan LK, Gera M, Choudhuri S, Lyle N, Ali KA, Diamond G. 2022. Antifungals and drug resistance. Encyclopedia 2:1722–1737. DOI: 10.3390/encyclopedia2040118.

29. Huber A, Hajdu D, Bratschun-Khan D, Gáspári Z, Varbanov M, Philippot S, Fizil Á, Czajlik A, Kele Z, Sonderegger C, Galgóczy L, Bodor A, Marx F, Batta G. 2018. New antimicrobial potential and structural properties of PAFB: A cationic, cysteine-rich protein from *Penicillium chrysogenum* Q176. Sci Rep 8:1751. DOI: 10.1038/s41598-018-20002-2.

30. Jacobsen ID. 2014. *Galleria mellonella* as a model host to study virulence of *Candida*. Virulence 5:237–239. DOI: 10.4161/viru.27434.

31. Kainz K, Bauer MA, Madeo F, Carmona-Gutierrez D. 202. Fungal infections in humans: the silent crisis. Microb Cell 7:143–145. DOI: 10.15698/mic2020.06.718.

32. Katiyar SK, Edlind TD. 2009. Role for Fks1 in the intrinsic echinocandin resistance of *Fusarium solani* as evidenced by hybrid expression in *Saccharomyces cerevisiae*. Antimicrob Agents Chemother 53:1772–1778. DOI: 10.1128/AAC.00020-09.

33. Kelly SL, Lamb DC, Kelly DE, Manning NJ, Loeffler J, Hebart H, Schumacher U, Einsele H. 1997. Resistance to fluconazole and cross-resistance to amphotericin B in *Candida albicans* from AIDS patients caused by defective sterol delta5,6-desaturation. FEBS Lett 400:80–82. DOI: 10.1016/s0014-5793(96)01360-9.

34. Klengel T, Liang WJ, Chaloupka J, Ruoff C, Schröppel K, Naglik JR, Eckert SE, Mogensen EG, Haynes K, Tuite MF, Levin LR, Buck J, Mühlschlegel FA. 2005. Fungal adenylyl cyclase integrates CO_2_ sensing with cAMP signaling and virulence. Curr Biol 15:2021–2026. DOI: 10.1016/j.cub.2005.10.040.

35. Kovács R, Holzknecht J, Hargitai Z, Papp C, Farkas A, Borics A, Tóth L, Váradi G, Tóth GK, Kovács I, Dubrac S, Majoros L, Marx F, Galgóczy L. 2019. *In vivo* applicability of *Neosartorya fischeri* antifungal protein 2 (NFAP2) in treatment of vulvovaginal candidiasis. Antimicrob Agents Chemother 63:e01777–18. DOI: 10.1128/AAC.01777-18.

36. Kovács R, Nagy F, Tóth Z, Forgács L, Tóth L, Váradi G, Tóth GK, Vadászi K, Borman AM, Majoros L, Galgóczy L. 2021. The *Neosartorya fischeri* antifungal protein 2 (NFAP2): A new potential weapon against multidrug-resistant *Candida auris* biofilms. Int J Mol Sci 22:771. DOI: 10.3390/ijms22020771.

37. Ksiezopolska E, Gabaldón T. 2018. Evolutionary emergence of drug resistance in *Candida* opportunistic pathogens. Genes 9:461. DOI: 10.3390/genes9090461.

38. LaFayette SL, Collins C, Zaas AK, Schell WA, Betancourt-Quiroz M, Gunatilaka AA, Perfect JR, Cowen LE. 2010. PKC signaling regulates drug resistance of the fungal pathogen *Candida albicans* via circuitry comprised of Mkc1, calcineurin, and Hsp90. PLoS Pathog 6:e1001069. DOI: 10.1371/journal.ppat.1001069.

39. Lemmon SK, Jones EW. 1987. Clathrin requirement for normal growth of yeast. Science 238:504–509. DOI: 10.1126/science.3116672.

40. Liang SH, Bennett RJ. 2019. The Impact of Gene Dosage and Heterozygosity on The Diploid Pathobiont *Candida albicans*. J Fungi (Basel). 6(1):10. DOI: 10.3390/jof6010010.

41. Lu Y, Su C, Ray S, Yuan Y, Liu H. 2019. CO_2_ signaling through the Ptc2-Ssn3 axis governs sustained hyphal development of *Candida albicans* by reducing Ume6 phosphorylation and degradation. mBio 10:e02320–18. DOI: 10.1128/mBio.02320-18.

42. Majoros WH, Pertea M, Salzberg SL. 2004. TigrScan and GlimmerHMM: two open source ab initio eukaryotic gene-finders. Bioinformatics 20:2878–2879. DOI: 10.1093/bioinformatics/bth315.

43. McColl AI, Bleackley MR, Anderson MA, Lowe RGT. 2018. Resistance to the plant defensin NaD1 features modifications to the cell wall and osmo-regulation pathways of yeast. Front Microbiol 9:1648. DOI: 10.3389/fmicb.2018.01648.

44. McMahon HT, Boucrot E. 2011. Molecular mechanism and physiological functions of clathrin-mediated endocytosis. Nat Rev Mol Cell Biol 12:517–533. DOI: 10.1038/nrm3151.

45. Morales-Johansson H, Jenoe P, Cooke FT, Hall MN. 2004. Negative regulation of phosphatidylinositol 4, 5-bisphosphate levels by the INP51-associated proteins TAX4 and IRS4. J Biol Chem 279:39604–39610. DOI: 10.1074/jbc.M405589200.

46. Morschhäuser J. 2016. The development of fluconazole resistance in *Candida albicans* - an example of microevolution of a fungal pathogen. J Microbiol 54:192–201. DOI: 10.1007/s12275-016-5628-4.

47. Nurk S, Bankevich A, Antipov D, Gurevich AA, Korobeynikov A, Lapidus A, Prjibelski AD, Pyshkin A, Sirotkin A, Sirotkin Y, Stepanauskas R, Clingenpeel SR, Woyke T, McLean JS, Lasken R, Tesler G, Alekseyev MA, Pevzner PA. 2013. Assembling single-cell genomes and mini-metagenomes from chimeric MDA products. J Comput Biol 20:714–737. DOI: 10.1089/cmb.2013.0084.

48. Papp C, Bohner F, Kocsis K, Varga M, Szekeres A, Bodai L, Willis JR, Gabaldón T, Tóth R, Nosanchuk JD, Vágvölgyi C, Gácser A. 2020 Triazole evolution of *Candida parapsilosis* results in cross-resistance to other antifungal drugs, influences stress responses, and alters virulence in an antifungal drug-dependent manner. mSphere 5:e00821–20. DOI: 10.1128/mSphere.

49. Pappas PG, Kauffman CA, Andes DR, Clancy CJ, Marr KA, Ostrosky-Zeichner L, Reboli AC, Schuster MG, Vazquez JA, Walsh TJ, Zaoutis TE, Sobel JD. 2016. Clinical practice guideline for the management of candidiasis: 2016 update by the infectious diseases society of America. Clin Infect Dis 62:e1–50. DOI: 10.1093/cid/civ933.

50. Payne GS, Hasson TB, Hasson MS, Schekman R. 1987. Genetic and biochemical characterization of clathrin-deficient *Saccharomyces cerevisiae*. Mol Cell Biol 7(11):3888– 3898. DOI: 10.1128/mcb.7.11.3888-3898.1987.

51. Perlin DS, Rautemaa-Richardson R, Alastruey-Izquierdo A. 2017. The global problem of antifungal resistance: prevalence, mechanisms, and management. Lancet Infect Dis 17:e383– e392. DOI: 10.1016/S1473-3099(17)30316-X.

52. Poissy J, Rouzé A, Cornu M, Nseir S, Sendid B. 2022. The changing landscape of invasive fungal infections in ICUs: A need for risk stratification to better target antifungal drugs and the threat of resistance. J Fungi (Basel) 8:946. DOI: 10.3390/jof8090946.

53. Popp C, Hampe IAI, Hertlein T, Ohlsen K, Rogers PD, Morschhäuser J. 2017. Competitive fitness of fluconazole-resistant clinical *Candida albicans* strains. Antimicrob Agents Chemother 61:e00584–17. DOI: 10.1128/AAC.00584-17.

54. Prasad T, Saini P, Gaur NA, Vishwakarma RA, Khan LA, Haq QM, Prasad R. 2005. Functional analysis of CaIPT1, a sphingolipid biosynthetic gene involved in multidrug resistance and morphogenesis of *Candida albicans*. Antimicrob Agents Chemother 49:3442– 3452. DOI: 10.1128/AAC.49.8.3442-3452.2005.

55. Rashid S, Correia-Mesquita TO, Godoy P, Omran RP, Whiteway M. 2022. SAGA Complex subunits in *Candida albicans* differentially regulate filamentation, invasiveness, and biofilm formation. Front Cell Infect Microbiol 12:764711. DOI: 10.3389/fcimb.2022.764711.

56. Rauseo AM, Coler-Reilly A, Larson L, Spec A. 2020. Hope on the horizon: Novel fungal treatments in development. Open Forum Infect Dis 7:ofaa016. DOI: 10.1093/ofid/ofaa016.

57. Rautemaa-Richardson R, Richardson MD. 2017. Systemic fungal infections. Medicine 45: 757–762. DOI: 10.1016/j.mpmed.2017.09.007.

58. Rollenhagen C, Agyeman H, Eszterhas S, Lee SA. 2022. *Candida albicans* END3 mediates endocytosis and has subsequent roles in cell wall integrity, morphological switching, and tissue invasion. Microbiol Spectr 10:e0188021. DOI: 10.1128/spectrum.01880-21.

59. Scott NE, Edwin Erayil S, Kline SE, Selmecki A. 2023. Rapid evolution of multidrug resistance in a *Candida lusitaniae* infection during micafungin monotherapy. Antimicrob Agents Chemother 67:e0054323. DOI: 10.1128/aac.00543-23.

60. Silva LV, Sanguinetti M, Vandeputte P, Torelli R, Rochat B, Sanglard D. 2013. Milbemycins: more than efflux inhibitors for fungal pathogens. Antimicrob Agents Chemother 57:873–586. DOI: 10.1128/AAC.02040-12.

61. Singh SD, Robbins N, Zaas AK, Schell WA, Perfect JR, Cowen LE. 2009. Hsp90 governs echinocandin resistance in the pathogenic yeast *Candida albicans* via calcineurin. PLoS Pathog 5:e1000532. DOI: 10.1371/journal.ppat.1000532.

62. Sionov E, Chang YC, Garraffo HM, Dolan MA, Ghannoum MA, Kwon-Chung KJ. 2012. Identification of a *Cryptococcus neoformans* cytochrome P450 lanosterol 14α-demethylase (Erg11) residue critical for differential susceptibility between fluconazole/voriconazole and itraconazole/posaconazole. Antimicrob Agents Chemother 56:1162–1169. DOI: 10.1128/AAC.05502-11.

63. Sonderegger C, Fizil Á, Burtscher L, Hajdu D, Muñoz A, Gáspári Z, Read ND, Batta G, Marx F. 2017. D19S mutation of the cationic, cysteine-rich protein PAF: Novel insights into its structural dynamics, thermal unfolding and antifungal function. PLoS One 12:e0169920. DOI: 10.1371/journal.pone.0169920.S

64. Sonderegger C, Galgóczy L, Garrigues S, Fizil Á, Borics A, Manzanares P, Hegedüs N, Huber A, Marcos JF, Batta G, Marx F. 2016. A Penicillium chrysogenum-based expression system for the production of small, cysteine-rich antifungal proteins for structural and functional analyses. Microb Cell Fact 15(1):192. DOI: 10.1186/s12934-016-0586-4

65. Sonderegger C, Váradi G, Galgóczy L, Kocsubé S, Posch W, Borics A, Dubrac S, Tóth GK, Wilflingseder D, Marx F. 2018. The evolutionary conserved γ-core motif influences the anti- *Candida* activity of the *Penicillium chrysogenum* antifungal protein PAF. Front Microbiol 9:1655. DOI: 10.3389/fmicb.2018.01655.

66. Sorgo AG, Heilmann CJ, Dekker HL, Bekker M, Brul S, de Koster CG, de Koning LJ, Klis FM. 2011. Effects of fluconazole on the secretome, the wall proteome, and wall integrity of the clinical fungus *Candida albicans*. Eukaryot Cell 10:1071–1081. DOI: 10.1128/EC.05011-11.

67. Tan PK, Davis NG, Sprague GF, Payne GS. 1993. Clathrin facilitates the internalization of seven transmembrane segment receptors for mating pheromones in yeast. J Cell Biol 123:1707–1716. DOI: 10.1083/jcb.123.6.1707.

68. Tóth L, Kele Z, Borics A, Nagy LG, Váradi G, Virágh M, Takó M, Vágvölgyi C, Galgóczy L. 2016. NFAP2, a novel cysteine-rich anti-yeast protein from *Neosartorya fischeri* NRRL 181: isolation and characterization. AMB Express 6:75. DOI: 10.1186/s13568-016-0250-8.

69. Tóth L, Váradi G, Borics A, Batta G, Kele Z, Vendrinszky Á, Tóth R, Ficze H, Tóth GK, Vágvölgyi C, Marx F, Galgóczy L. 2018. Anti-candidal activity and functional mapping of recombinant and synthetic *Neosartorya fischeri* antifungal protein 2 (NFAP2). Front Microbiol 9:393. DOI: 10.3389/fmicb.2018.00393.

70. Van Daele R, Spriet I, Wauters J, Maertens J, Mercier T, Van Hecke S, Brüggemann R. 2019. Antifungal drugs: What brings the future? Med Mycol 57:S328–S343. DOI: 10.1093/mmy/myz012.

71. Wang J, Chitsaz F, Derbyshire MK, Gonzales NR, Gwadz M, Lu S, Marchler GH, Song JS, Thanki N, Yamashita RA, Yang M, Zhang D, Zheng C, Lanczycki CJ, Marchler-Bauer A. 2023. The conserved domain database in 2023. Nucleic Acids Res. 51(D1):D384–D388. DOI: 10.1093/nar/gkac1096.

72. White T, Pfaller M, Rinaldi M, Smith J, Redding S. 1997. Stable azole drug resistance associated with a substrain of *Candida albicans* from an HIV-infected patient. Oral Diseases 3: S102–S109. DOI: 10.1111/j.1601-0825.1997.tb00336.x

73. World Health Organization. 2022. WHO fungal priority pathogens list to guide research, development and public health action. ISBN: 978-92-4-006024-1 Licence: CC BY-NC-SA3.0 IGO.

74. Yeater KM, Chandra J, Cheng G, Mukherjee PK, Zhao X, Rodriguez-Zas SL, Kwast KE, Ghannoum MA, Hoyer LL. 2007. Temporal analysis of *Candida albicans* gene expression during biofilm development. Microbiology 153:2373–2385. DOI: 10.1007/s11046-016-0088-2.

