## Supplementary materials for "The *Neosartorya (Aspergillus) fischeri* antifungal protein NFAP2 has low potential to trigger resistance development in *Candida albicans in vitro*"

**TABLE S1** Positions of mutations in the amino acid sequences of the affected proteins in *Neosartorya* (*Aspergillus*) *fischeri* antifungal protein 2 (NFAP2/4, 5, 6) and fluconazole (FLC/3, 5, 6) resistant strains.

| **CHC1p** in FLC/3 | |
| --- | --- |
| Amino acid sequence | MSNDIPIDFTELTQLTELGIPQTSLDFKSTTLESDHYICVRESGAQGNTVAIVNLKNNNEVTRKNMTADNAIMHPKEFVISLRANGTTLQIFNLGSKQRLKAFTMDQPVIFWKWLDDTYLGLVTQSSIYYWNVFDGTNNGPTKLTDRHHTLNNCQIINFVAEPDLNWFAVTGIAQEDGRIAGHIQLYSKSRNVSQAIEGHVCKFASISLSGGVQPTKVFCVGNKNAQGQGNMHIIEIDHVDGNPQFQKKVVDIFFPPDASNDFPISLQASNKYGIVYVLTKYGFIHLYDMETGANLFVNRITADPVFTASSYNDGTGLITINKAGQVLSVEVSRDKIIPYVLDKLSNVPLALALSSRGGFPGAENLFQQQFQTYLNQGDYSNAAKVAASSEQLRTQDTINKLKNITPQPGQISPILQYFSTLLDRGTLNKFESIELAKPVLQQDRKPLFEKWLKEDKLTSSEELGDIVKSYNDTTLAL**A(*)V(I)**YIRANVNIKVVSCLAELGQFDKILPYCQKVGYNPDYTNLIQNLVRVNPDKASEFATSLLSSPDANLNVEQIADLFFSQNYIQQGTAFLLDALKNDTPAEGHLQTKVLEINLLHAPQVADAILGNQMFSHYDKPTIGKLCEKSGLFQRALEHYDDLKDIKRVIVHTNVLPSDWLVSYFGQLNVDQSVACIKELLSNNMQQNLQVVIQVATKYSDLIGAAKLIKIFEEYKCTEGLYYYLSSIVNLTQDPDVVFKYIQAAARMNQTKEIERVVRDNNVYNGEKVKNFLKEFKLDDQLPLIIVCDRFNFVHDLILYLYKNQYFKFIEVYVQSVNPANTPQVVAGLLDVDCDENIIKGLLMSVLGRVPIKELVEEVEKRNRLKILLPFLEKTLEGGSNDQEVYNTLAKIYIDSNNSPEKFLQENNNYDTLVVGKYCEKRDPYLAYISYSKGGNDDQLISITNENKMYKYQARYLLSKSDIELWNKVLSSDNIHRRQLVDQVISTGIPELDDPEPISITVKAFMENDLPEELMELLEKIILEPSPFSENASLQGLMILTAIKADSSKVSSYIEKLDKFDPNEIAPLCIDNGLNEEAFEVYDKFELRPEAMKVLVEDIVSLDRAEQYAEKYDTPELWYQLGTAQLDGLRIPEAIDSYVKSKNPENFAQVIEIAEHAGKEEELITFLDMARETLREPVVDGALINAYATLDRLSDMEKFVGGSNVADLEAIGDKLFEAKNYKGAKILYSNVSKYSKLATTLVYLGDYQGAVDCARKASNTQVWKQVNSACIENKEFRLAQICGLNLIIDAEELPELVKTYEYNGYFNELIALFENGLSLERAHMGMFTELAILYAKYSPEKVMEHLKLFWSRINIPKVLTACEDAHLYPELIFLYCHYEEWDNAALTMIEKSEVAFDHSSFKEIIVKAPNLEIHYKAIQFYMNENPSLLVDLLKVLTPKLDLPRVVRMFVQSDNLPMIKPFLISVLDKNNSVVNSAYHDLLIEEEDYKSLRSSIENESNNRFNKLDLAERLEKHDLIFFRQIAATLYTKEKKFNRAISILKTDKLWPDLLRTVAISKSKKIAHELLDYFVETGNHECFVALLYTSYEFIANDYVMELSWLHNLSNFIKPYEISIAFENQKKLNEVYQDLQKRKEADRKQEEEPGVGQPLMLTNGPMS YQGTGATGIGYQPTGTGFGNAF* |
| **INP51p** in FLC/5 | |
| Amino acid sequence | MKLSTIFTAFAATIATVAGYETTGSKQTVDILIDYIIKETPELSQNDVANWENGDTVTLQYVVNNNEESEITVVGVTGQFKNPVNNEIVTNLTTGKVGPIAVPPGEAIKFDQKINVDLIPANYELIPHVFIAQDSLIKVIPCRGQLATIVDAAVSFFDPRLIFLELVLLITFAGLIYVGYEIWGKQYFKGVAPVKAKKVSAAKASSPVASGPSTTSATGYDTNWIPESHLKQKKTKKGLVSGHSKDKDQNKDTKVLVEFVLKEYLDLSLYRDITPKHGGLLGLLGLLNVKGKTFIGFITRDEWTASATVTDRIYKITDTEFYCINNDEYDYLLDKEYENMSHQERERLRYPAASVQRLLSSGAFYYSKQFDMTSNIQERGFVSSDYKLIADSSFFKSFMWNGFMTEELIETRKRMSPAEQKIIDKSGLLIIVIRGYAKTVNTTVGGCEALMTLISKQSCAKEGPLFGDWGSDGDGYVSNYLESEIIIYTEKFCLSYVIVRGNVPMYWELENNFSTKTILAANGKQIAFPRSFEASQEALVRHFDRLSSQYGDIHVLNALSDKSYKGVLNSAYEEQLKYFLQNRESTDIGYKVLYTRIPIASSRIKKIGYSGQNPYDIVSLLSNSIIDFGALFYDSKPNSFIGKQLGVFRINSFDSLNKANFLSKIISQEVIDLAFRDIGLELDRELYVKHAQLWEENDLWISKLTLNFASTSDKLHTSHNSIKSSFVKSHITKKYFGGVVESKPNEIAMLKLLGRLQDQSPVTMFNPIHNYVNKELNKRAKDFTSKLDLSVYASTFNVNGSVYEGDIDKWIYPEENDYDLIFVGLQEIVVLNAGQMVNTDFRNKTQWERKILGVLQKRNKYMVMWSGQLGGVALYFFVKESQVKYVSNVECSFKKTGL**G(V)**GVSANKGGIAVSFKFSDTTICFVSAHLAAGLSNIEERHQNYKALIKGIQFSKNRRIQNHDAVIWLGDFNYRIDLTNDQELNEEKDASLPAPSSDKHKWWLEGGKAAKIIIPGLEDDNMVMNPWRPINPFEKSNEPEFVSKNDLEAIQN* |
| **PGA58p** in NFAP2/5 and NFAP2/6 | |
| Amino acid sequence | MYCLAAAPSKVGTLDGPVELETEEAGCSGTEAGADGAGAGEDSGCSGAGADGAGAGADSGCSGAGKAGADGAGADGAGAGTDSGCSGAGGAGADGAGAGVDSASFVVGAAGCSGCSGAGAGADGAGAGAAGEAGCSGAGADGSGAAGVAGVAGAAGCSGVAGEAGCSGAGADGSGAAGVAGVAGVAGCSGCSGAGADGSG**V(A)**AGVAGAAGVAGAAGVAGVAGCSGCSGAGADGSGAAGVAGVAGCSGCSGAGADGAGAGADGSGCSGAGGAGAEAEGAGAGVAGEEAGQFVCSGPHEVMVTTVEDSTV* |
| **PTC2p** in FLC/3 | |
| Amino acid sequence | MEDAHATILNLYDLPLKKSLSSNSEQDEDEEDEEEEADSQDSTTTNTTKSQQQQNQKEDLQQNNDKQNLQQDIKDGDNDTQMSDSDENHHHAAPQQQHIAFFGVYDGHGGEKAAIFTGEKLHHLIKETKEFKQKDYINALKQGFLNCDQEILKDFYMRDDDSGCAATSAIITPDLIVCGNAGDSRTIMSTNGFAKALSFDHKPSNEGEKARICAAGGYVDMGRVNGNLALSRGIGDFDFKKNVDLPAEEQIVTCYPDQQQKNDPNSLNKYGPISQPYNELYKEMYGEYYDIGQQGQGGNGRSNIGSGSGSGRGSGASGSSSGGGASINGNSIFGRSGYDDEEDDDEEKNIERRRRDQDESNLNGGAISLQKLLASNAITNENGVIYLDTSSAQSLLAHFGVADAGAHGDEEDEEEEGEGEDVHQGLAHEEEEEEE**E(*)**E**E(*)**EGSKIEEISEKIDDEV* |
| **RIM101p** in FLC/6 | |
| Amino acid sequence | MNYNIHPVTYLNADSNTGASESTASHHGSKKSPSSDIDVDNATSPSSFTSSQSPHINAMGNSPHSSFTSQSAANSPITDAKQHLVKPTTTKPAAFAPSANQSNTTAPQSYTQPAQQLPTQLHPSLNQAYNNQPSYYLHQPTYGYQQQQQHQEFNQPSQQYHDHHGYYSNNNILNQNQPAPQQNPVKPFKKTYKKIRDEDLKGPFKCLWSNCNIIFETPEILYDHLCDDHVGRKSSNNLSLTCLWENCGTTTVKRDHITSHLRVHVPLKPFHCDLCPKSFKRPQDLKKHSKTHAEDHPKKLKKAQRELMKQQQKEAKQQQKLANKRANSMNATTASDLQLNYYSGNPADGLNYDDTSRKRRYENNSQHNMYVVNSILNDFNFQQMAQAPQQPGVVGTAGSAEFTTKRMKAGTEYNIDVFNKLNHLDDHLHHHHPQQQHPQQQYGGNIYEAEKFFNSLSNSIDMQYQNMSTQYQQQHAGSTFAQQKPTQQASGQLYPSLPTIGNGSYTTSGSSHKEGLVNNHNGYLPSYPQINRSLPYSSGVAQQPPSALEFGGVSTYQKSAQLYEEDSSDSSEEDD**Y(C)**STSSEDELDTLFDKLNIDDNKVEEVTIDGFNLKDVAKHREMIHAVLGYLRNQIEQQEKEKSKEQKEVDVNETKLYPTITAF* |
| **UBIPp** in NFAP2/4 | |
| Amino acid sequence | MSEFPNIDTLDFIPAAAGTGTGTGVAQQQQQQRQFQQYLRTPKSASSPKFQLTATNFSNQTTPAFGSVATFSNNSFLPDGSDSDLLPANAQFTPITPGIVNYNTYMATQSVQPNNKFEPLLSTPNSVNGVSNSPVQFSPPTQQF**S(Ø)**PPTQQYPPSSSSSSSAPASTVGSTVGEKTGATIGLGVSIEPAVKHKKQPSPHQQAALHFQQQLNNEAAAKTHKHSGGSSTTKTRTQLSPEMQKLIEQQREKQQQQLLLLQKGEIPKPPELKPKRKSSCRPHKQKRRYIPEEEQYYQHQYQHQHRRIASLPLQTIPGDSPTTIEVSSPQLEVCNDNSFGCNTTSFITGGNITAEELLSNDPFEVLDDHRNNIFPEFVDLDFEDIGGVDFENSIESVDVLSLNKNSSSTNVSVQRPTLKKKKSITPDPAKIVKTLKKASSFSNGSGISGSSSPTFIINPKFQWTKSQTTEQQQRLLKKKVPYQHAHSLPDLSIQEMPVFTLSNHYLFVYENADSILQDENIEENSQTSPRKQQQLQQLQEQQSNKEVELLQNLESGLVEFKLELNKNKMS* |

The amino acids in the mutation positions are highlighted by colored bold letters and in grey. The native amino acids are followed by the mutant variants indicated in parentheses. Amino acids are categorized by color based on their distinctive properties: Orange: small nonpolar, green: hydrophobic, magenta: polar, red: positively charged, blue: negatively charged. CHC1: clathrin heavy chain 1, INP51: phosphoinositide 5-phosphatase, PGA58: putative glycosylphosphatidylinositol-anchored protein, PTC2: serine/threonine-specific protein phosphatase, RIM101: alkaline-responsive transcriptional regulator, UBIP: Uncharacterized biofilm-induced protein, *: termination codon, Ø: deletion.

**TABLE S2** Oligonucleotides and reaction parameters of PCR applied for amplification of the mutated gene regions in the consequence of NFAP2 or FLC pressure in a microevolution experiment.

| **CHC1** | | | |
| --- | --- | --- | --- |
| Primer A: | CHC1_m_fw (5'-3') | GTAATGCCGCTAAAGTTGCC | Tm: 57.3 C° |
| Primer B: | CHC1_m_rev (5'-3') | CAAGTTGGCATCTGGACTTG | Tm: 57.3 C° |
|  | Reaction parameters | Initial denaturation of genomic DNA was carried out at 98 °C for 5 minutes, then 30 cycles of the following: denaturation at 98 °C for 20 seconds, annealing at 55 °C for 20 seconds, and extension at 72 °C for 20 seconds. Final extension was carried out at 72 °C for 5 minutes. | |
| **INP51** | | | |
| Primer A: | INP51_m_fw (5'-3') | GTTGAATAAGCGTGCCAAGG | Tm: 57.3 C° |
| Primer B: | INP51_m_rev (5'-3') | GGTCGCCATGGATTCATAAC | Tm: 57.3 C° |
|  | Reaction parameters | Initial denaturation of genomic DNA was carried out at 98 °C for 5 minutes, then 30 cycles of the following: denaturation at 98 °C for 20 seconds, annealing at 55 °C for 20 seconds, and extension at 72 °C for 30 seconds. Final extension was carried out at 72 °C for 5 minutes. | |
| **PGA58** | | | |
| Primer A: | PGA58_m_fw2 (5'-3') | AGCTGGTTGTTCTGGAACTG | Tm: 57.3 C° |
| Primer B: | PGA58_m_fw3 (5'-3') | TGTTTAGCAGCAGCACCTTC | Tm: 57.3 C° |
| Primer C: | PGA58_m_rev2 (5'-3') | ATCTGCTCCAGCTCCTGAAC | Tm: 59.4 C° |
|  | Reaction parameters | Initial denaturation of genomic DNA was carried out at 98 °C for 5 minutes, then 30 cycles of the following: denaturation at 98 °C for 20 seconds, annealing at 56 °C for 20 seconds, and extension at 72 °C for 25 seconds. Final extension was carried out at 72 °C for 5 minutes. | |
| **PTC2** | | | |
| Primer A: | PTC2_m_fw1 (5'-3') | CAAGGTCAAGGTGGAAATGG | Tm: 57.3 C° |
| Primer B: | PTC2_m_rev2 (5'-3') | TCTTCAATCTTGGATCCTTC | Tm: 53.2 C° |
|  | Reaction parameters | Initial denaturation of genomic DNA was carried out at 98 °C for 5 minutes, then 30 cycles of the following: denaturation at 98 °C for 20 seconds, annealing at 53 °C for 20 seconds, and extension at 72 °C for 25 seconds. Final extension was carried out at 72 °C for 5 minutes. | |
| **RIM101** | | | |
| Primer A: | RIM101_m_fw (5'-3') | GCAAGTGGCCAATTGTATCC | Tm: 57.3 C° |
| Primer B: | RIM101_m_rev (5'-3') | TCTCTGTGCTTGGCAACATC | Tm: 57.3 C° |
|  | Reaction parameters | Initial denaturation of genomic DNA was carried out at 98 °C for 5 minutes, then 30 cycles of the following: denaturation at 98 °C for 20 seconds, annealing at 55 °C for 20 seconds, and extension at 72 °C for 15 seconds. Final extension was carried out at 72 °C for 5 minutes. | |
| **UBIP** | | | |
| Primer A: | Unchar_m_fw (5'-3') | TACTGGTGTGGCTCAACAAC | Tm: 57.3 C° |
| Primer B: | Unchar_m_rev (5'-3') | CTACCGCCACTGTGCTTATG | Tm: 59.4 C° |
|  | Reaction parameters | Initial denaturation of genomic DNA was carried out at 98 °C for 5 minutes, then 30 cycles of the following: denaturation at 98 °C for 20 seconds, annealing at 56 °C for 20 seconds, and extension at 72 °C for 20 seconds. Final extension was carried out at 72 °C for 5 minutes. | |

PCR were carried out using Phusion High-Fidelity DNA Polymerase with 5×Phusion HF Buffer (Thermo Fischer Scientific, Waltham, MA, USA) according to the manufacturer’s instructions. The PCR mixture contained 1×Phusion HF Buffer 200 µM each dNTP, 0.5 primer µM primer A, 0.5 primer µM primer B, 0.02 U/µl Phusion High-Fidelity DNA Polymerase, 100 ng genomic DNA. CHC1: clathrin heavy chain 1, INP51: phosphoinositide 5-phosphatase, PGA58: putative glycosylphosphatidylinositol-anchored protein, PTC2: serine/threonine-specific protein phosphatase, RIM101: alkaline-responsive transcriptional regulator, UBIP: Uncharacterized biofilm-induced protein.

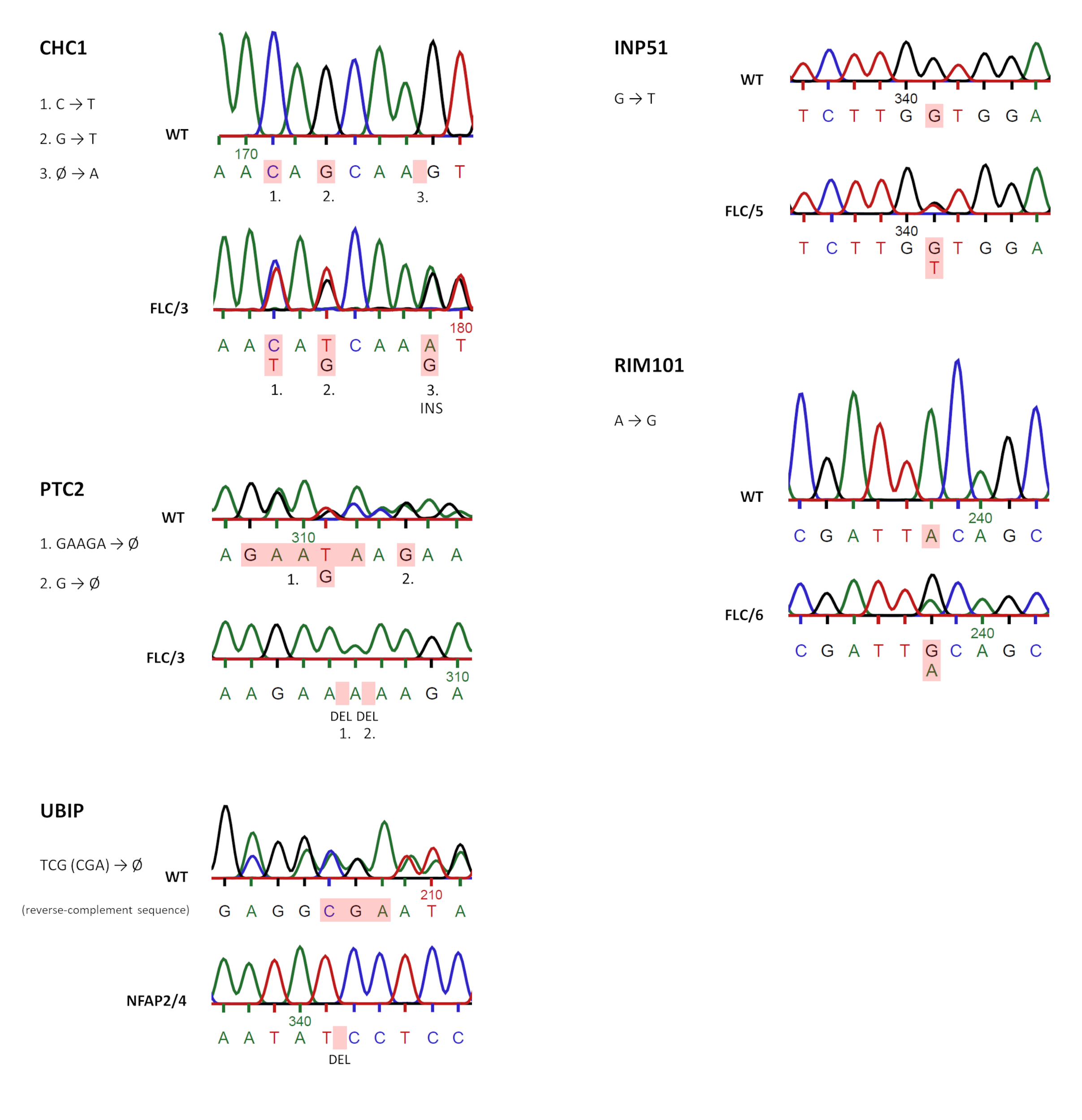

**FIGURE S1** Sequenograms of the mutated gene regions in the consequence of *Neosartorya* (*Aspergillus*) *fischeri* NRRL 181 antifungal protein 2 (NFAP2) or fluconazole (FLC) pressure in a microevolution experiment in comparison with the related regions of the wild-type (WT) *Candida albicans* CBS 5982 genes. Base changes are highlighted in red. Double peaks in the mutated region indicate heterozygous mutations. CHC1: clathrin heavy chain 1, DEL: deletion, FLC/3, 5, 6: FLC-resistant strains, INP51: phosphoinositide 5-phosphatase, INS: insertion, NFAP2/4, 5, 6: NFAP2-resistant strains, PTC2: serine/threonine-specific protein phosphatase, RIM101: alkaline-responsive transcriptional regulator, UBIP: Uncharacterized biofilm-induced protein. The full length sequenograms files can be found in Supplementary File 2.

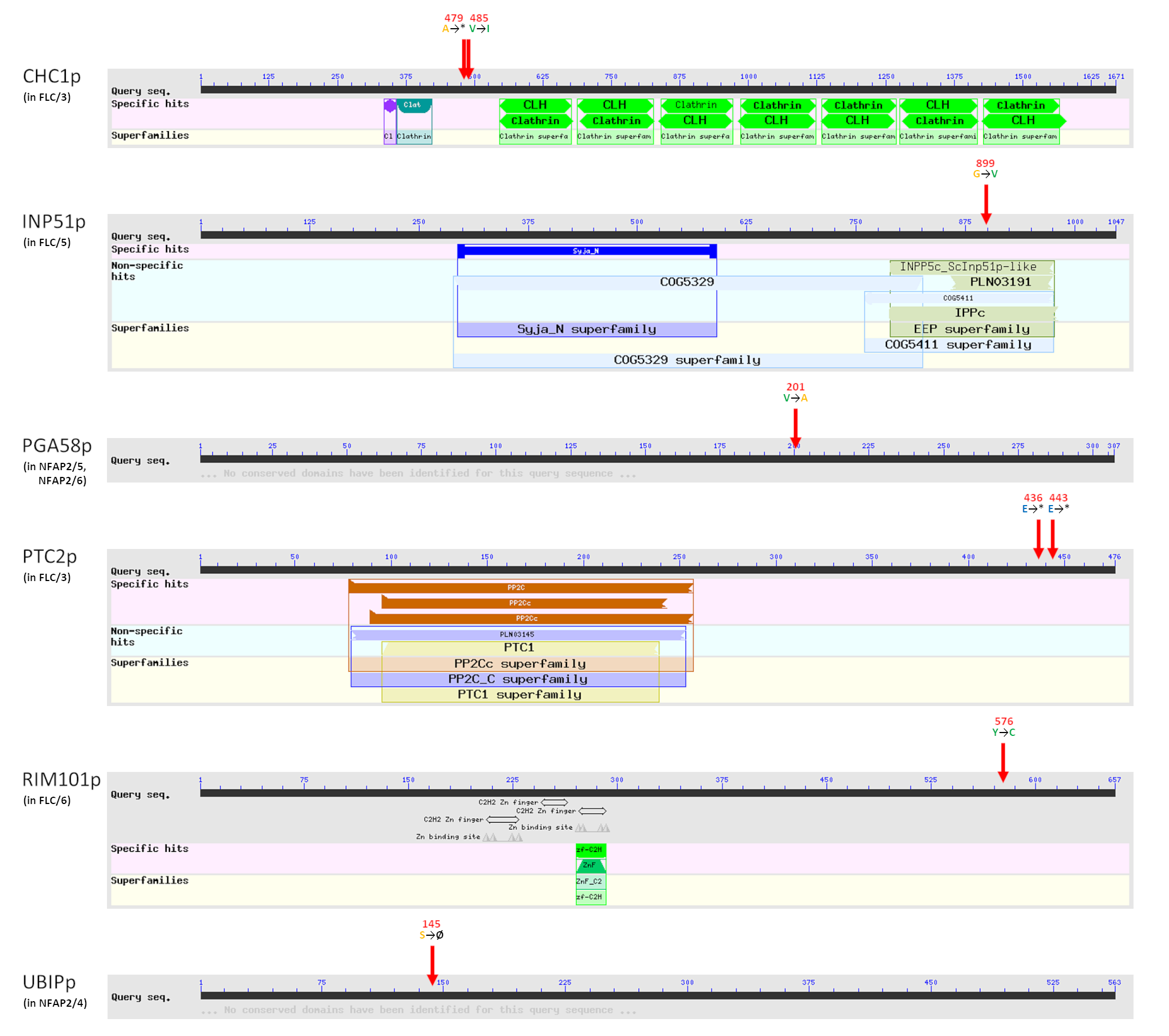

**FIG S2** Locations of detected *Candida albicans* CBS 5982 gene mutations at amino acid level as consequences of *Neosartorya* (*Aspergillus*) *fischeri* NRRL 181 antifungal protein 2 (NFAP2) or fluconazole (FLC) pressure. Conserved domains are indicated below query seq. The specific positions (amino acid number) of respective mutations in the sequence are indicated with red arrows. The amino acid alterations are indicated with colored letters. Amino acids are categorized by color based on their distinctive properties: Orange small nonpolar, green: hydrophobic, magenta: polar, red: positively charged, blue: negatively charged. CHC1: clathrin heavy chain 1, INP51: phosphoinositide 5-phosphatase, PGA58: putative glycosylphosphatidylinositol-anchored protein, PTC2: serine/threonine-specific protein phosphatase, RIM101: alkaline-responsive transcriptional regulator, UBIP: Uncharacterized biofilm-induced protein, *: termination codon, Ø: deletion. Graphical summaries were generated by Conserved Domain Search (https://www.ncbi.nlm.nih.gov/Structure/cdd/wrpsb.cgi; default options with “Standard display” result mode) using Conserved Domain Database (https://www.ncbi.nlm.nih.gov/cdd/).

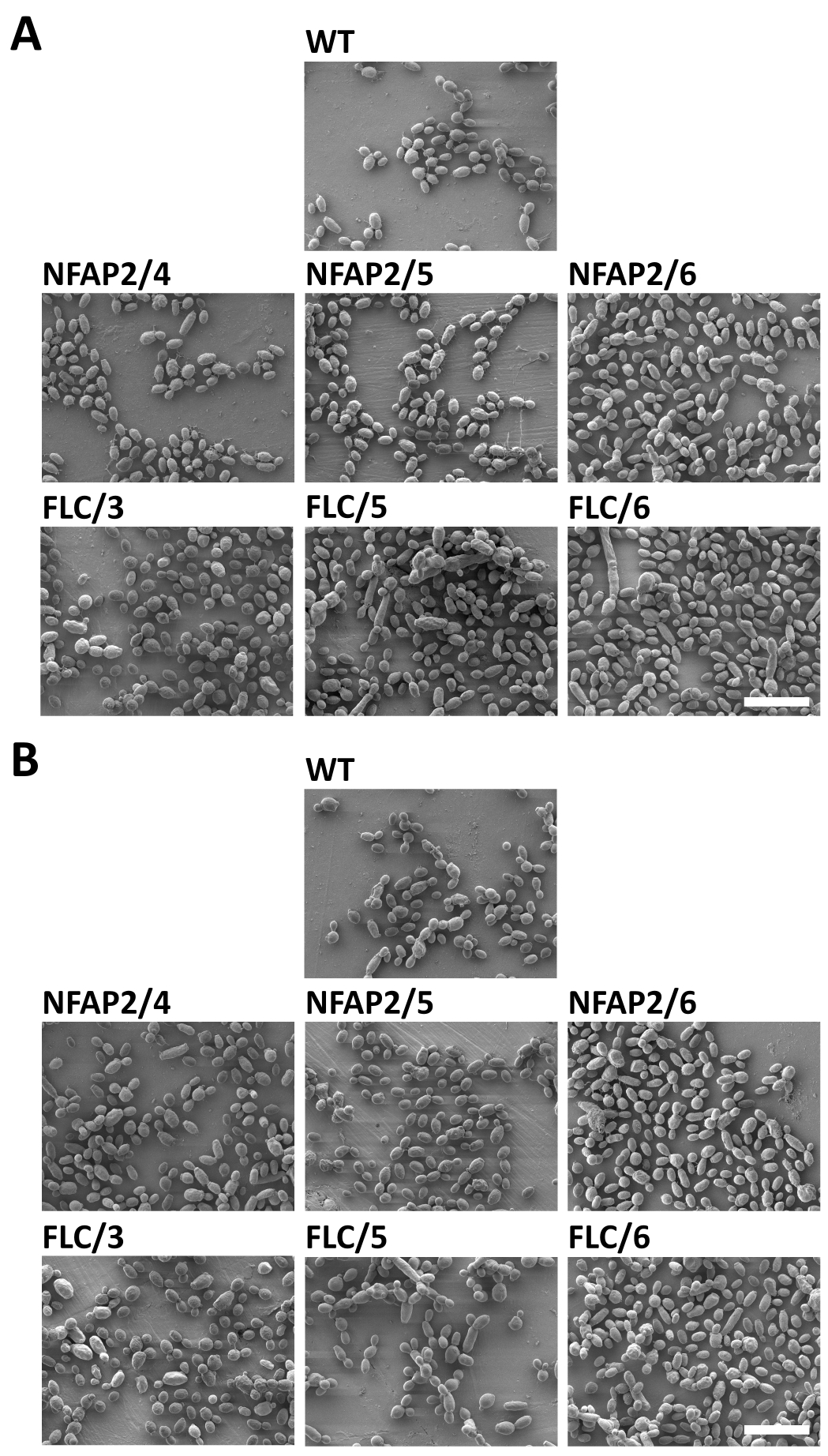

**FIG S3** Scanning electron microscopy of the parental wild-type *Candida albicans* CBS 5982 (WT), *Neosartorya* (*Aspergillus*) *fischeri* antifungal protein 2- resistant (NFAP2/4, 5, 6), and fluconazole-resistant (FLC/3, 5, 6) strains (A), and the impact of NFAP2-treatment (1 × minimum inhibitory concentration, 30 min, 30°C, 160 rpm, low-cation medium) on cell morphology (B). 2,000× magnification. Scale bars represent 10 µm.

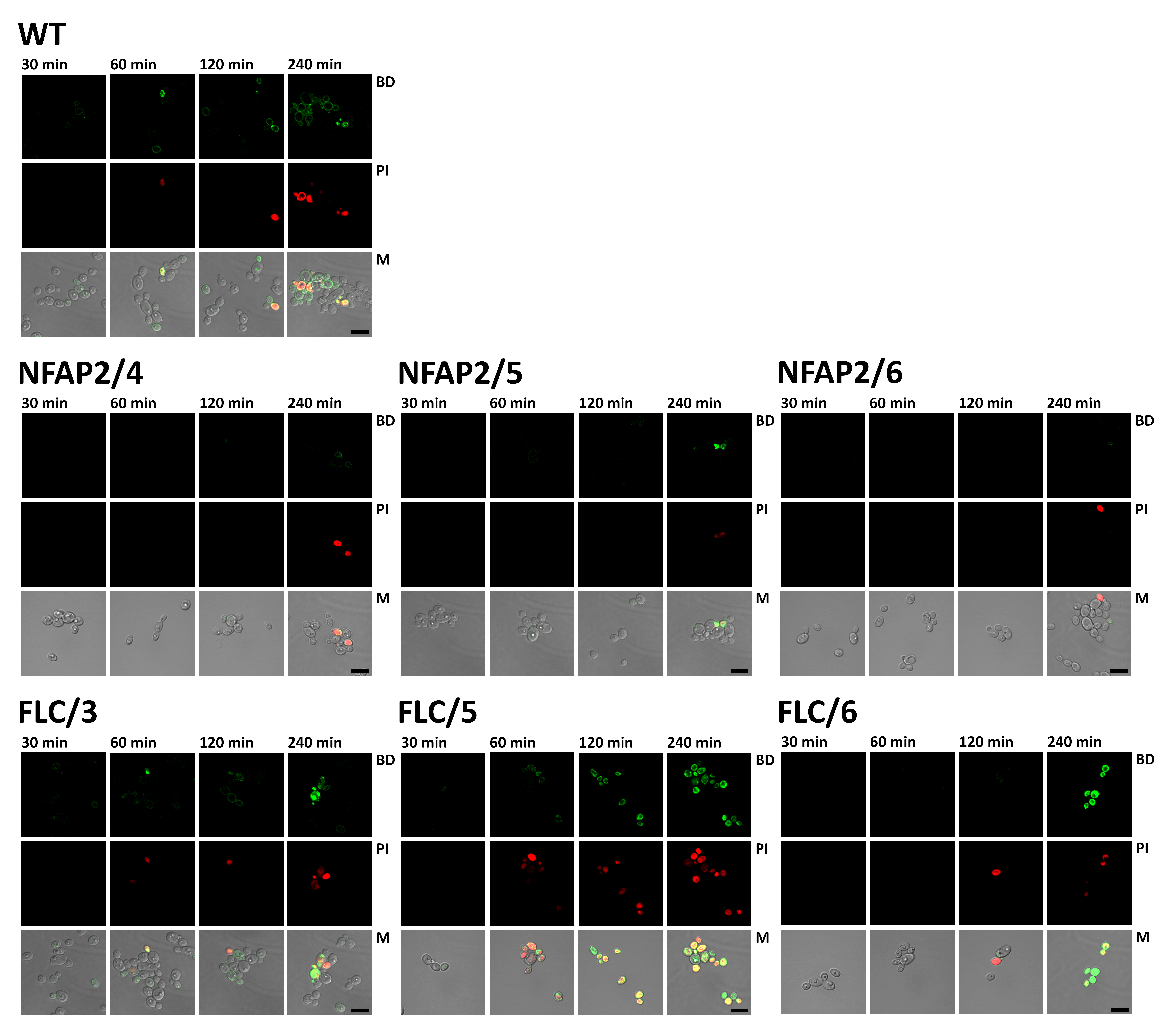

**FIG S4** Confocal laser scanning microscopy of the parental wild-type CBS 5982 (WT), NFAP2-resistant (NFAP2/4, 5, 6), and fluconazole-resistant (FLC/3, 5, 6) *Candida albicans* CBS 5982 strains (4 × 10^6^ cells) after treatment with the green fluorophore BODIPY-labelled NFAP2 (Bd-NFAP2, 3.125 µg ml^−1^) for 30, 60, 120, and 240 minutes (30°C, 160 rpm, low cationic medium), and co-stained with propidium-iodide (PI) for 10 minutes. Sequential scanning was done for Bd-NFAP2 (BD, green) and PI (PI, red) at 488 and 543 nm, respectively. M: merged BP, PI, and bright field pictures. Scale bars represent 10 µm. The gamma settings of the merged images have been adjusted to uniformize the tones of the backgrounds for illustrative purposes.

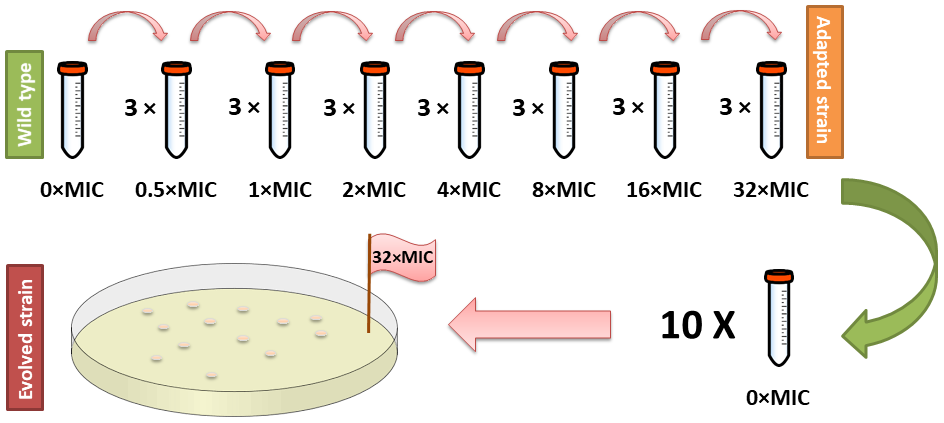

**FIG S5** Schematic representation of microevolution experiment performed in low cationic medium to generate NFAP2- or fluconazole-resistant *Candida albicans* strains.
